# Environmental and molecular control of tissue-specific ionocyte differentiation in zebrafish

**DOI:** 10.1101/2024.01.12.575421

**Authors:** Julia Peloggia, Mark E. Lush, Ya-Yin Tsai, Christopher Wood, Tatjana Piotrowski

## Abstract

Organisms adjust their physiology to cope with environmental fluctuations and maintain fitness. These adaptations occur via genetic changes over multiple generations or through acclimation, a set of reversible phenotypic changes that confer resilience to the individual. Aquatic organisms are subject to dramatic seasonal fluctuations in water salinity, which can affect the function of lateral line mechanosensory hair cells. To maintain hair cell function when salinity decreases, ion-regulating cells, Neuromast-associated ionocytes (Nm ionocytes), increase in number and invade lateral line neuromasts. How environmental changes trigger this adaptive differentiation of Nm ionocytes and how these cells are specified is still unknown. Here, we identify Nm ionocyte progenitors as *foxi3a/foxi3b*-expressing skin cells and show that their differentiation is associated with sequential activation of different Notch pathway components, which control ionocyte survival. We demonstrate that new Nm ionocytes are rapidly specified by absolute salinity levels, independently of stress response pathways. We further show that Nm ionocyte differentiation is selectively triggered by depletion of specific ions, such as Ca^2+^ and Na^+^/Cl^−^, but not by low K^+^ levels, and is independent of media osmolarity. Finally, we demonstrate that hair cell activity plays a role in Nm ionocyte recruitment and that systemic factors are not necessary for Nm ionocyte induction. In summary, we have identified how environmental changes activate a signaling cascade that triggers basal skin cell progenitors to differentiate into Nm ionocytes and invade lateral line organs. This adaptive behavior is an example of physiological plasticity that may prove essential for survival in changing climates.

## INTRODUCTION

Organisms have evolved various strategies to thrive in fast-changing environments. One crucial mechanism that enhances their adaptability is phenotypic plasticity, where the same genotype leads to different phenotypes in an environment-dependent manner^1,2^. Many examples of phenotypic plasticity have been studied in detail, such as the adaptation of organisms to different temperatures, body size regulation in sea urchins, fur coat changes of arctic foxes, hypoxia tolerance in fishes and responses of organisms to osmotic stresses^2–6^.

In mammals and other terrestrial vertebrates, osmoregulation, which is the control of water and ion concentrations in the body, occurs mainly through dietary intake and urine production in the kidney^7^. While this is a systemic regulation, multiple tissues have additional mechanisms to fine-tune the control of the ionic balance and osmoregulation. Their ion composition is maintained in part by tissue-specific ion-regulating cells named ionocytes. Ionocytes express ion channels that actively transport ions across the cell membranes and establish proper ionic concentration and osmotic pressures^6,8^. In mammals, locally acting tissue-specific ion-regulating cells have been identified in the ear, epididymis, kidney and lungs, and loss of function of ion channels highly expressed in these cells leads to hearing loss, infertility, renal tubular acidosis and cystic fibrosis, respectively^9–13^.

Aquatic animals, particularly those living in freshwater habitats, are subjected to much greater fluctuations in environmental salinity and pH^14^. Consequently, these organisms require fast and robust adaptation mechanisms to thrive in these dynamic conditions. To control ion homeostasis and osmoregulation, teleost fishes also rely on ionocytes^4,6^. To date, five main ionocyte subtypes have been identified in zebrafish and are classified based on their expression of different channels and their specific roles in regulating ions: HR (H^+^ secretion/Na^+^ uptake/NH4^+^ excretion), NaR (Ca^2+^ uptake), NCC (Na^+^/Cl^−^ uptake), KS (K^+^ secretion) and SLC26 ionocytes (Cl^−^ uptake/ HCO3^−^ secretion)^15^. While ionocytes were first described in the eel gill epithelium^16^, it is now appreciated that they also associate with organs such as the embryonic skin of fish and frogs and the zebrafish sensory lateral line^17,18^.

The zebrafish lateral line is a mechanosensory organ composed of multiple sensory units, called neuromasts, distributed along the trunk and head (Figure 1A and 1B). These neuromasts contain sensory hair cells that detect water flow and help with orientation, prey detection and predator avoidance^19^. We previously demonstrated that upon changes in salinity and pH, basal skin cells migrate, invade mature lateral line neuromasts and differentiate into neuromast-associated ionocytes (Nm ionocytes, Figure 1B and 1C)^18^. We named this process *adaptive cell invasion* and showed that Nm ionocytes help fine tune the activity of hair cells, ensuring their functionality across a wide range of environmental conditions. However, how environmental changes are detected and what signals are required to specify Nm ionocytes are still not understood.

**Figure 1:**
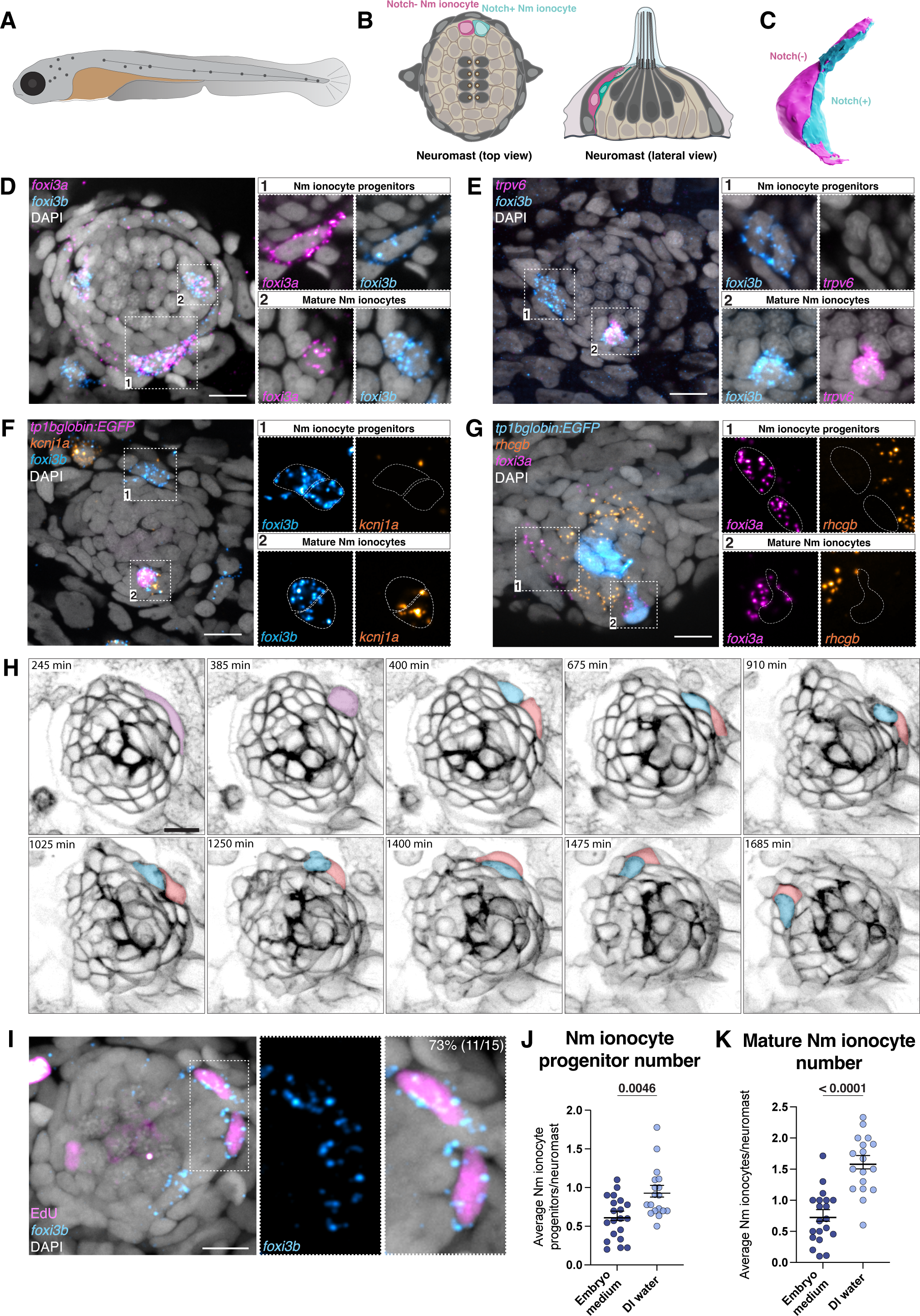
Nm ionocyte progenitors are *foxi3a/foxi3b*-positive cells derived from one cell division event. (A) Schematic of a zebrafish larva at 5 dpf. Lateral line neuromasts are labeled in dark grey. (B) Schematic of a neuromast top view (left) and lateral view (right). Nm ionocyte pair is labeled in magenta and cyan. (C) 3D modeling of Nm ionocyte pair. (D) HCR labeling of transcription factors *foxi3a* and *foxi3b* shows two mature Nm ionocytes inside a lateral line neuromast (2) and a ventrally located double-positive progenitor adjacent to the neuromast (1). (E) Maximum intensity projection of HCR for *foxi3b* and *trpv6* labeling a pair of progenitors and one of mature Nm ionocytes. (F) Maximum intensity projection of HCR for *foxi3a* and *kcnj1a* in the Notch reporter (*tp1bglobin:EGFP*) background. (G) Maximum intensity projection of *foxi3a* and *rhcgb* HCRs in the Notch reporter (*tp1bglobin:EGFP*) background. (H) Maximum intensity projection of still images of a time-lapse of *cldnb:lyn-EGFP* fish at 3 dpf. A pair of cells invades a neuromast. (I) EdU incorportation in combination with *foxi3b* HCR (J) Average number of Nm ionocyte progenitors per neuromast for embryos incubated in embryo medium (n = 20 larvae) or in DI water for 48h (n = 18 larvae; Mann-Whitney test). (K) Average number of mature Nm ionocyte per neuromast for embryos incubated in embryo medium or in DI water for 48h (Unpaired t-test). All scale bars = 10 µm.

Here, we identify for the first time the progenitors that give rise to mature Nm ionocytes and show that their survival is dependent on Notch signaling. We demonstrate that Nm ionocyte specification is triggered by absolute levels of salinity and that the salinity sensor discriminates between different ions. Finally, we show that environmental changes do not depend on systemic responses to specify new Nm ionocytes and that hair cell function plays a role in this process. Collectively, we elucidate the earliest cellular and molecular events required for Nm ionocyte specification and differentiation and shed light on the mechanisms required for sensory organ adaptation to different environmental changes.

## RESULTS

### Identification of Nm ionocyte progenitors

We previously showed that decreased salinity and pH lead to the adaptive differentiation of basal stem cells into a pair of Nm ionocytes, which invade lateral line neuromasts^18^. How zebrafish detect environmental changes and activate this adaptive response is still unknown. Here, we sought to identify the early steps of ionocyte induction and differentiation. The transcription factors *foxi3a* and *foxi3b* are major regulators of zebrafish skin and Nm ionocyte development, and their loss inhibits ionocyte differentiation^18,20,21^. To test if *foxi3a/3b* mRNAs label Nm ionocyte progenitors before invasion we performed whole mount hybridization chain reaction RNA fluorescent *in situ* hybridization (HCR). We observed cells immediately adjacent to neuromasts that co-express both *foxi3a/3b* (Figure 1D, inset 1). Inside the neuromast, *foxi3a* is expressed only in the Notch-negative Nm ionocyte, while *foxi3b* is expressed in both cells of the mature Nm ionocyte pair (Figure 1D, inset 2). This is consistent with the molecular similarities of these cells with HR and NaR skin ionocytes, respectively (Figure S1A-B)^18^. These data indicate that *foxi3a/foxi3b*-expressing cells adjacent to neuromasts could be the Nm ionocyte progenitors.

To further investigate the identity and differentiation state of the cells while outside neuromasts, we performed HCRs for genes expressed in differentiated ionocytes in combination with a Notch reporter (*tp1bglobin:EGFP*) that labels one cell of the mature Nm ionocyte pair. The Ca^2+^ channel *trpv6* is expressed in skin and mature Nm ionocytes but not in the putative progenitors outside the neuromasts (Figure 1E and S1C). Likewise, the ion channels *kcnj1a.1, rhcgb* and *slc4a1b* are only expressed in differentiated ionocytes (Figure 1F-G and S1D-F). These findings suggest that the cells outside the neuromasts are not fully differentiated or functional and could be the progenitors that give rise to Nm ionocytes. Additionally, these cells are crescent shaped and morphologically distinct from mature skin ionocytes, which are bigger, with round nuclei and square cell morphology (Figure S1C-H). These morphological differences further support the notion that these cells are not a subtype of differentiated skin ionocytes, but Nm ionocyte progenitors.

The fact that mature Nm ionocytes are always present in pairs implies that they might arise by cell division of a single progenitor or stem cell. To test if Nm ionocyte progenitors divide we performed an ethynyl-2’-deoxyuridine (EdU) incorporation assay followed by *foxi3b* HCR. We observed that the majority of Nm ionocyte progenitors were EdU-positive, and often appeared in pairs (Figure 1I). Time-lapse analyses of Nm ionocyte development also showed that a single precursor divides and that the resulting pair invades neuromasts (Figure 1H and Supplementary Video 1), confirming that the pair of mature Nm ionocytes arises by cell division of a single progenitor cell. Taken together, these data show that Nm ionocyte progenitors are pairs of *foxi3a/foxi3b*-positive cells that derive from a single cell division event.

### Nm ionocyte progenitors are induced adjacent to neuromasts by low salinity

Nm ionocyte differentiation is an adaptive behavior triggered by changes in media salinity and pH^18^. We next tested if the number of *foxi3a/foxi3b*-positive Nm ionocyte progenitors is dependent on salinity. Incubation of zebrafish in deionized water (DI water) for 48 hours increases the total progenitor number per neuromast (average number), as well as the number of neuromasts with Nm ionocyte progenitors (frequency) (Figure 1J and S1K). Consistent with the higher progenitor number, the average number and frequency of mature Nm ionocytes also increase following 48h of salinity decrease (Figure 1K and S1L), similar to our earlier findings^18^. The increase in Nm ionocyte progenitor frequency in low salinity suggests that new progenitors are induced in response to environmental stimuli. Additional analyses of *foxi3a* and *foxi3b* HCRs show the presence of a subset of cells surrounding the neuromasts that express only *foxi3a*, while few *foxi3b*-only cells were detected (Figure S1I and S1J). This suggests that, like in skin ionocytes, *foxi3a* is activated before *foxi3b*^18,20,21^. Moreover, we detected individual *foxi3a*- and *foxi3a/b*-positive cells, which indicates these transcription factors are upregulated before cell division (Figure 1D and S1I). We conclude from this data that new Nm ionocyte progenitors are induced adjacent to neuromasts by low salinity and that *foxi3a* and *foxi3b* are upregulated prior to cell division.

### Components of the Notch signaling pathway are expressed in a precise spatio-temporal pattern during Nm ionocyte differentiation

Once inside neuromasts, each cell of the progenitor pair adopts a specific ionocyte fate, shown by the differential expression of transcription factors and ion channels^18^. To address the molecular basis of Nm ionocyte fate determination, we focused on the Notch signaling pathway, as its expression has been observed in Nm ionocytes. Notch signaling is an ancestral pathway that controls cell identity and cell fate decisions via the interaction of a ligand with a receptor expressed in a neighboring cell^22^. Their interaction results in the cleavage of the Notch receptor intracellular domain (NICD), which translocates to the nucleus and drives changes in gene expression^23,24^. Zebrafish possess four receptors and nine different Notch ligands (five Deltas and four Jaggeds), and interactions of Notch receptors with different ligands lead to distinct downstream outputs^25^.

One of the two mature Nm ionocytes expresses the receptor *notch1b* (Figure 2A)^18^. To test if other Notch receptors are present in mature Nm ionocytes and progenitors, we performed HCRs of *notch1a*, *notch1b* and *notch3* in combination with a Notch reporter (*tp1bglobin:EGFP*) or *foxi3b* HCR, respectively. *notch1b* is expressed in both *foxi3b*-positive progenitors (Figure 2B), while *notch1a* and *notch3* are neither expressed in progenitors nor in mature Nm ionocytes (Figure S2A-E). This contrasts with the skin, where *notch1a* and *notch3*, but not *notch1b*, play a role in ionocyte specification^20,21^. To determine which ligands are expressed in mature Nm ionocytes, we performed HCR for all known Delta and Jagged ligands. We detected expression of only *dll4* in the Notch-negative Nm ionocyte, and no other ligands in the mature pair (Figure 2C and S2F). Thus, mature Nm ionocyte pairs express a *notch1b*-*dll4* receptor-ligand combination. To determine if Notch signaling could be involved in specifying distinct fates in the progenitors, we analyzed expression of ligands by HCR before invasion of neuromasts. We observed expression of *dll4* in both progenitor cells, as well as of a second ligand, *dld* (Figure 2D and Figure S2G). However, not all *foxi3b*-positive progenitors expressed these two Notch ligands. To determine the dynamics of *dll4* and *dld* expression in both progenitors, we analyzed their frequency in combination with *foxi3b*. We observed a fraction of progenitors that only express *foxi3b*, suggesting it is transcribed before Notch ligands are expressed (Figure 2E). A small percentage of progenitors co-express either *foxi3b/dld* or *foxi3b/dll4*, and almost half of the progenitors express all three genes (*foxi3b, dld* and *dll4*). The most parsimonious interpretation is that *dld* is expressed prior to *dll4* and is downregulated before neuromast invasion.

**Figure 2:**
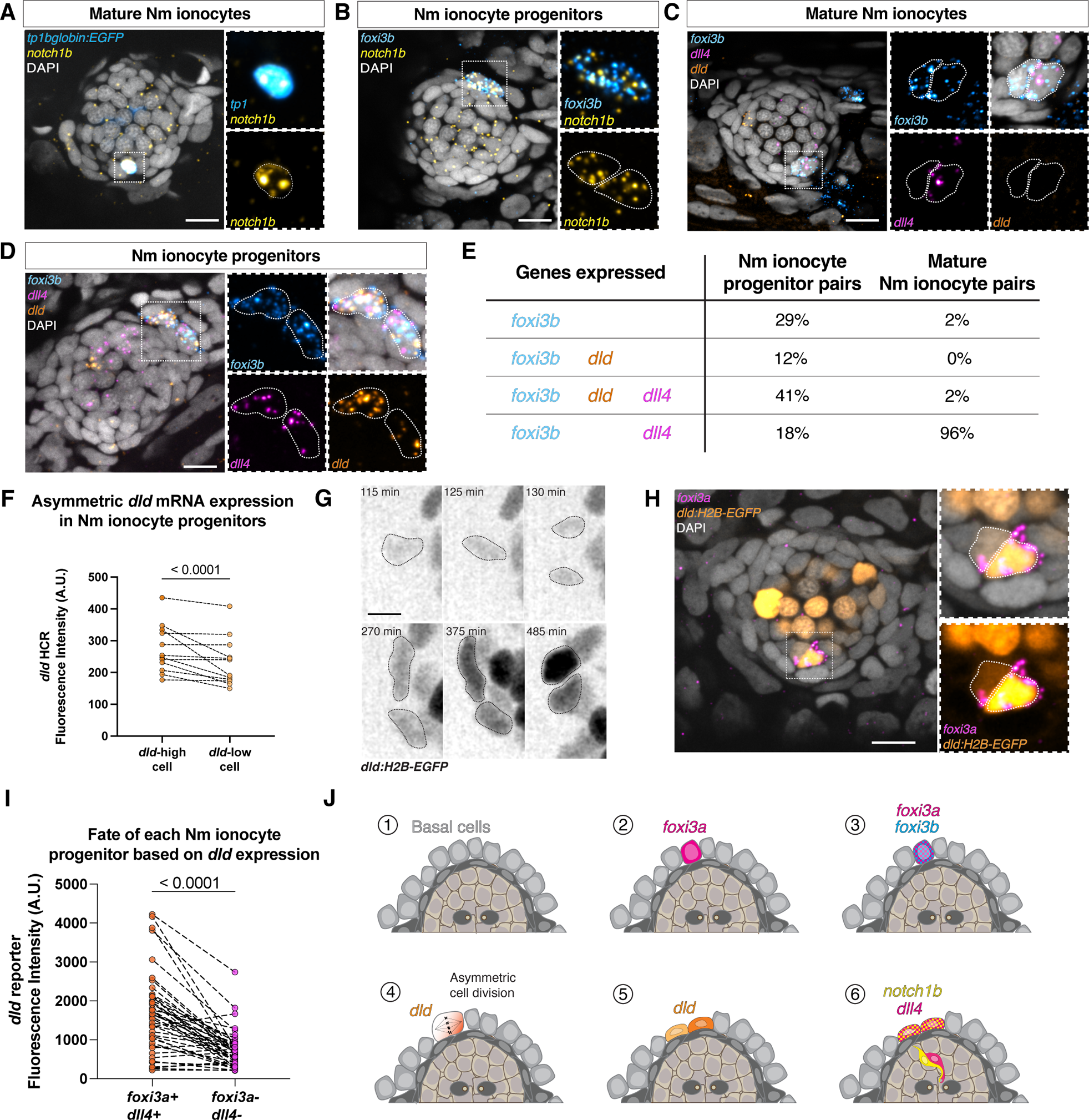
A switch of Notch ligand expression is associated with ionocyte differentiation and invasion of lateral line organs in zebrafish. (A) Confocal image of *notch1b* HCRs in mature Nm ionocytes and (B) Nm ionocyte progenitors. (C) Representative image of triple HCR for *foxi3b, dll4* and *dld* expression in mature Nm ionocytes and (D) Nm ionocyte progenitors. (E) Summary table of percentages of Nm ionocyte pairs with expression of genes *foxi3b, dll4* and *dld* (n = 51 mature Nm ionocytes and 17 progenitor pairs). (F) Quantification of *dld* mRNA expression between the two cells of the progenitor pair (Unpaired t-test). (G) Maximum intensity projection of *Tg(dld:H2B-EGFP)^psi84Tg^* combined with HCR for *foxi3a*. The *dld* reporter labels both Nm ionocytes and lateral line hair cells (H) Quantification of EGFP fluorescence in each Nm ionocyte of the pair and its relation to *foxi3a* expression (n = 45 pairs of cells, 24 larvae; Wilcoxon matched-pairs signed rank test). (I) Time-lapse analysis of *dld:H2B-EGFP* shows *dld* is upregulated prior to cell division, scale bar = 5 µm. Gamma was changed in the *dld:H2B-EGFP* channel to allow for visualization of dim structures. (J) Summary of gene expression during Nm ionocyte development. Scale bars = 10 µm unless specified otherwise.

Our data suggest that the Notch ligand *dld* is transiently expressed in progenitors. Additionally, we noticed an asymmetry in the levels of *dld* expression in progenitors shown by HCR (Figure 2D and 2F). To investigate the dynamic expression of *dld* in more detail, we generated a reporter line by knocking-in H2B-EGFP into the 5’UTR of *dld* (*dld:H2B-EGFP*) with CRISPR-Cas12a. Time-lapse analyses of the *dld:H2B-EGFP* reporter show that *dld* is expressed prior to mitosis and that the expression of GFP becomes asymmetric after cell division (Figure 2G), resulting in a *dld:H2B-EGFP*^high^ and a *dld:H2B-EGFP*^low^ cell. The histone tagged GFP is stable, making Nm ionocytes fluorescent as they invade neuromasts and differentiate, and the GFP can therefore be used as a lineage tracer. HCR for *foxi3a* in *dld:H2B-EGFP* transgenic larvae shows that the *dld:H2B-EGFP*^high^ cell differentiates into the *foxi3a*-positive Nm ionocyte, which also expresses *dll4* (Figure 2H, 2I and S2F).

Altogether, our results demonstrate that different Notch pathway components are dynamically expressed during Nm ionocyte differentiation. *foxi3a* and *foxi3b* are expressed in the progenitor cells prior to cell division and before Notch signaling. Then, *dld* is transiently expressed in both daughter cells as they divide and is downregulated as the progenitors invade neuromasts and differentiate. Concomitant with the *dld* pulse, *dll4* and *notch1b* are upregulated in both progenitors but each gets restricted to one of the mature cells, with *dll4* being restricted to the *dld*^high^ and *notch1b* to the *dld*^low^ cell (Figure 2J).

Notch signaling culminates in the NICD-dependent transcriptional activation of several genes, including genes in the *hairy/enhancer of split (her)* family^26^. These transcription factors repress genes in the Notch-positive cell that are activated in the Delta/Jagged-positive cell. We identified candidate downstream factors in our previously published single cell RNA sequencing data (scRNAseq) of ionocytes (Figure S3A) and performed HCRs. *hes6* and *her15* are expressed in the *notch1b*-positive mature ionocytes, and *her15* is expressed in some, but not all progenitors, while the remaining candidates are not detected in Nm ionocytes (Figure S3B-H). Additionally, *her15* expression is Nm ionocyte-specific, as it is not detected in skin ionocytes (Figure S3I and S3J). These results show that Notch signaling activates different downstream targets in the neuromast and skin ionocyte populations, and that *her15* is a specific target of *notch1b/dll4* signaling in Nm ionocytes.

### dll4 regulates ionocyte survival

To dissect the roles of the Notch ligands *dll4* and *dld* in Nm ionocyte specification and differentiation, we generated *dld* and *dll4* mutants using CRISPR/Cas12a (Figure S4A and S4E). Even though the *dld* mutant shows somitogenesis defects similar to other alleles^27^, the number and type of ionocytes do not change in the skin or neuromasts (Figure 3A-C and S3G-I). Likewise, *dld* overexpression driven by a heat shock promoter (*hsp70*) does not affect Nm ionocyte number (Figure S4J-M). We conclude that *dld* is not essential for Nm ionocyte determination and differentiation. In contrast, *dll4* mutants show a drastic reduction of Nm ionocytes both in embryo medium and DI water compared to wild-type siblings (Figure 3D and 3E). However, the average number of Nm ionocyte progenitors per fish does not change in *dll4* mutants (Figure 3F). Thus, *dll4* is required for development of mature Nm ionocytes, but not for progenitor specification.

**Figure 3:**
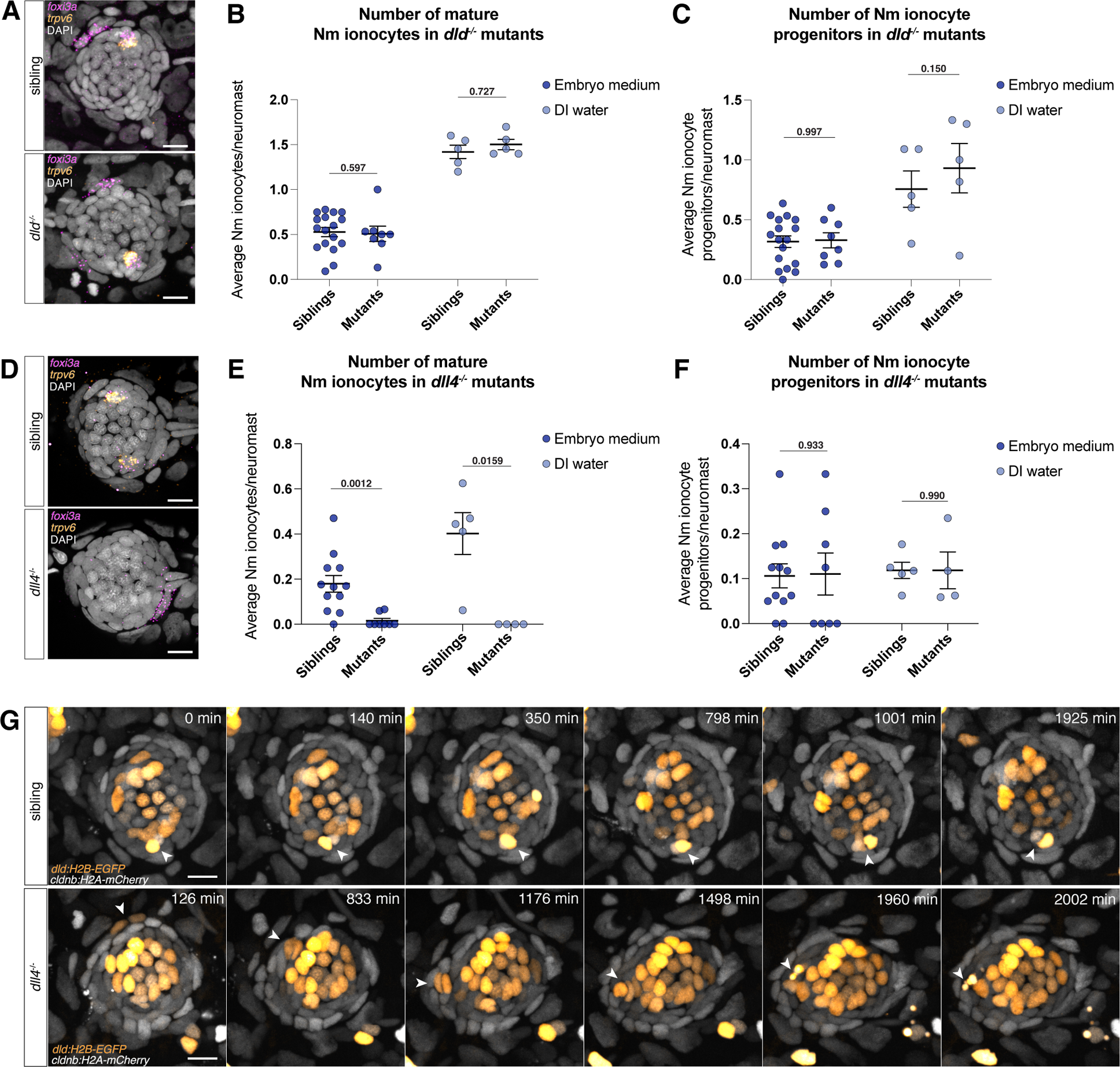
*dll4* affects Nm ionocyte differentiation but not progenitor specification. (A) Confocal images of neuromasts of wild-type siblings (top) and *dld* mutants (bottom) showing presence of mature Nm ionocytes and progenitors. (B) Number of mature Nm ionocytes and (C) progenitors in both siblings (embryo medium, n= 17 larvae, DI water, n = 5 larvae) and *dld* mutants embryo medium, n= 8 larvae, DI water, n = 5 larvae, unpaired t-test) raised in either embryo medium or incubated in DI water. (D) Confocal images of neuromasts of wild-type siblings (top) and *dll4* mutants (bottom) showing presence of mature Nm ionocytes and a progenitor, respectively. (E) Number of mature Nm ionocytes and (F) progenitors in both siblings (embryo medium, n= 12 larvae, DI water, n = 5 larvae) and *dll4* mutants (embryo medium, n= 8 larvae, DI water, n = 4 larvae, Mann-Whitney test) raised in different salinities. (G) Time-lapse still images showing a mature Nm ionocyte in siblings and (H) an invading Nm ionocyte pair in *dll4* mutants that undergoes cell death. Cells are labeled with *dld:H2B-EGFP* and *cldnb:H2A-mCherry*. Scale bars = 10 µm.

To determine why *dll4* mutants lack mature Nm ionocytes we performed time lapse analyses of mutants and siblings using the *dld:H2B-EGFP* reporter that labels Nm ionocytes and hair cells. We observed Nm ionocytes invading neuromasts in both siblings and mutants from 3 to 5 dpf. However, we observed a higher incidence of Nm ionocyte death in mutants compared to wild type larvae, indicating *dll4* plays a role in cell survival following invasion (Figure 3G, Supplementary Movies 2 and 3). This is consistent with our previous results which found that pharmacological inhibition of Notch signaling leads to death of mature ionocytes^18^. It is also possible, however, that the cell death phenotype we observe in *dll4* mutants is a consequence of aberrant differentiation. The lack of mature Nm ionocytes in *dll4* mutants prevents us from assessing the role of *dll4*-*notch1b* signaling in Nm ionocyte fate specification.

Previous work described that *dll4* is not expressed in the skin and it is not required for skin ionocyte differentiation^20,21^. However, we observed that while the total skin ionocyte (HR and NaR) density was not changed in 5 dpf *dll4* mutants (Figure S4B), the proportion of ionocyte subtypes was altered. *foxi3a*-positive HR ionocytes increase in density in the mutants, while *trpv6*-positive NaR skin ionocytes are totally absent in the skin of *dll4* mutants and are present only in the gills (Figure S4C and S4D). Indeed, a subset of skin ionocytes express *dll4* at 5 dpf (Figure S4F). Taken together, our results show that *dll4* regulates the proportion of HR and NaR skin ionocytes.

### New Nm ionocyte progenitors are rapidly induced by low salinity

It is not known how environmental stimuli are sensed at a cellular and molecular level and translated into progenitor activation and ionocyte differentiation. To determine the nature of the sensor, we first characterized Nm ionocyte differentiation dynamics upon salinity decrease, specifically the upregulation of *foxi3a/b* transcription factors. Time course analyses after DI water incubation showed that *foxi3a/b*-positive Nm ionocyte progenitors are quickly induced after salinity decrease, with a peak around 1 hour post incubation (Figure 4A-B). This rapid response suggests that the link between the sensor and the effector likely does not rely on detection by sensory systems, processing in higher brain centers and relayed through hormonal changes, but rather on less or faster components. Additionally, we observed that the number of progenitors oscillates over time, with a peak after DI water incubation and subsequent decrease around 4-8 hours after incubation, which likely correlates with these cells differentiating and invading the neuromasts. Progenitor oscillation is also observed in time-lapses, where new Nm ionocyte progenitors upregulate *dld:H2B-EGFP* and invade neuromasts, but new progenitors are not detected for several hours based on *dld:H2B-EGFP* expression (Supplementary Movie 2).

**Figure 4:**
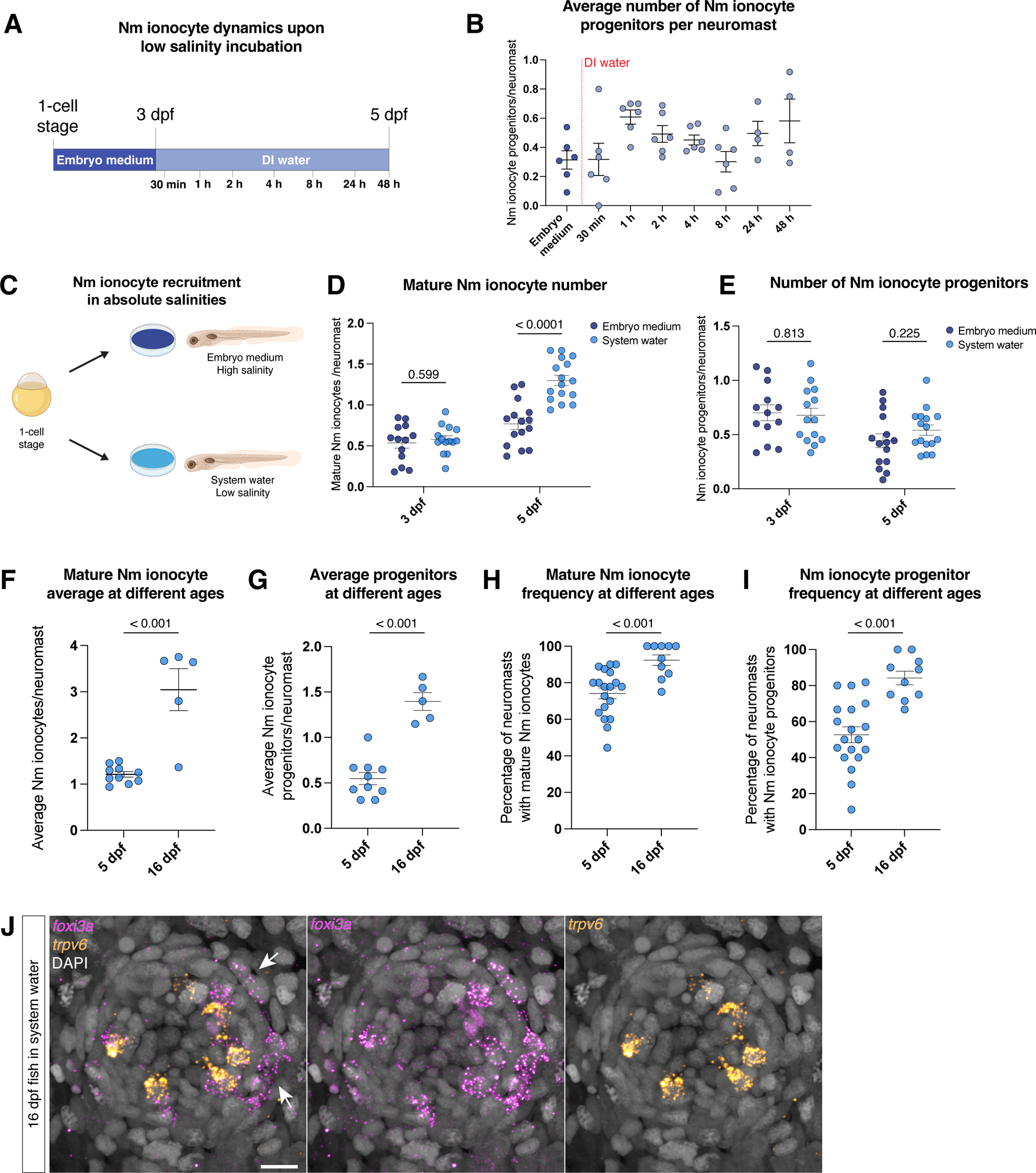
Nm ionocytes are rapidly induced by absolute low salinity. (A) Schematic representation of the time course experiment. (B) Average number of Nm ionocyte progenitors per neuromast following salinity decrease (C) Schematic representation of absolute salinity experiment. (D) Average number of mature ionocytes and (E) progenitors per neuromast of fish raised in different salinities at 3 and 5 dpf (Unpaired t-test). (F) Average mature ionocyte (G) and progenitor number in 5 dpf (n = 10 larvae) and 16 dpf (n = 5 larvae) larvae raised in system water (Unpaired t-test). (H) Frequency of mature and (I) Nm ionocyte progenitors at 5 and 16 dpf raised in system water (Unpaired t-test). (J) Representative HCR showing mature and Nm ionocyte progenitors in a neuromast of a 16 dpf fish.

A fast upregulation of Nm ionocyte progenitors could be caused by a stress response following salinity changes. To test if incubation in low salinity triggers stress response pathways, we tested 1) the activation of glucocorticoid (GR) signaling based on the expression of the downstream factor *dusp1*^28,29^, and 2) expression of components of the immediate early response AP-1 complex, *fosab* and *junba*. GR induces skin ionocyte proliferation upon salinity changes^30,31^, and AP-1 is activated in neuromasts upon hair cell death^32^. DI water incubation does not induce *dusp1* transcription in the skin nor in the neuromast (Figure S5A-C). We observed an increase in *fosab* mRNA expression 15 and 30 min after DI water incubation, and a transient slight increase of *junba* after 15 min incubation in DI water (Figure S5D-F). We next checked the expression of these genes in embryos incubated in system water. System water has an intermediate salinity compared to embryo medium and DI water and, consequently, the resulting frequency of Nm ionocytes. However, incubation in system water does not lead to expression of *fosab* and *junba* (Figure 4D and Figure S5D-F). This suggests that, while components of the AP-1 pathway are transcribed after DI water incubation, they may not be required for activating Nm ionocyte progenitors.

We next sought to test if Nm ionocyte induction requires salinity changes or if it is triggered by absolute levels of salinity. We incubated zebrafish embryos in different media starting at the one-cell stage and quantified Nm ionocyte numbers at 3 and 5 dpf (Figure 4C). At 3 dpf, numbers of mature Nm ionocytes and progenitors are not different between larvae raised in embryo medium (high salinity) or zebrafish system water (low salinity). However, at 5 dpf mature Nm ionocyte numbers in larvae raised in low salinity increase (Figure 4D), while progenitor Nm ionocyte numbers does not change (Figure 4E). Further, we do not see the activation of stress response pathways in constant low salinity (Figure S5G-I). These data demonstrate that the sensor that controls Nm ionocyte numbers detects absolute levels of salinity and not a change in salinity.

Mature Nm ionocytes increase in number from 3 to 5 dpf regardless of the salinity of the medium used. The number of progenitors, however, does not change within this time frame. This could be due to the progenitor turnover mentioned above. We previously demonstrated that the number of mature Nm ionocytes increases in adult animals^18^. To assess if progenitor numbers also change with age, we performed HCR for *foxi3a* and *trpv6* in 16 dpf juveniles. We observed that average mature Nm ionocyte number is higher in juveniles than in larvae, but the number of progenitors does not scale to the same proportions, with most neuromasts still having one or two pairs of progenitors but containing up to 6 mature Nm ionocyte pairs (Figure 4F and 4G). Meanwhile, the frequency of mature Nm ionocytes and progenitors is higher in juveniles when compared to 5 dpf larvae in the same salinity (Figure 4H-J). We conclude from this data that progenitors are likely a finite population around neuromasts and that the increase in adult mature Nm ionocyte numbers is due to increased invasion events.

### Differentiation of new Nm ionocytes is selectively triggered by different ions

Thus far, we determined that the salinity sensor responds fast, does not require a stress response and detects absolute salinity. We next asked which specific components of the medium are being detected, and incubated embryos for 48h in embryo media lacking specific ions. Incubation in medium lacking Ca^2+^ ions leads to an increase of mature Nm- and skin NaR ionocytes (Figure 5A and Figure S6). Additionally, lack of the majority of Na^+^ and Cl^−^ ions in the medium also leads to an increase in Nm ionocyte number, but to a lesser extent, even though Na^+^ and Cl^−^ ions are present in much higher concentrations in regular medium than Ca^2+^. Meanwhile, K^+^ depletion does not lead to an increase of Nm ionocytes. These results demonstrate that specific ions, and not general changes in medium ionic concentration, play a role in inducing new Nm ionocytes.

**Figure 5:**
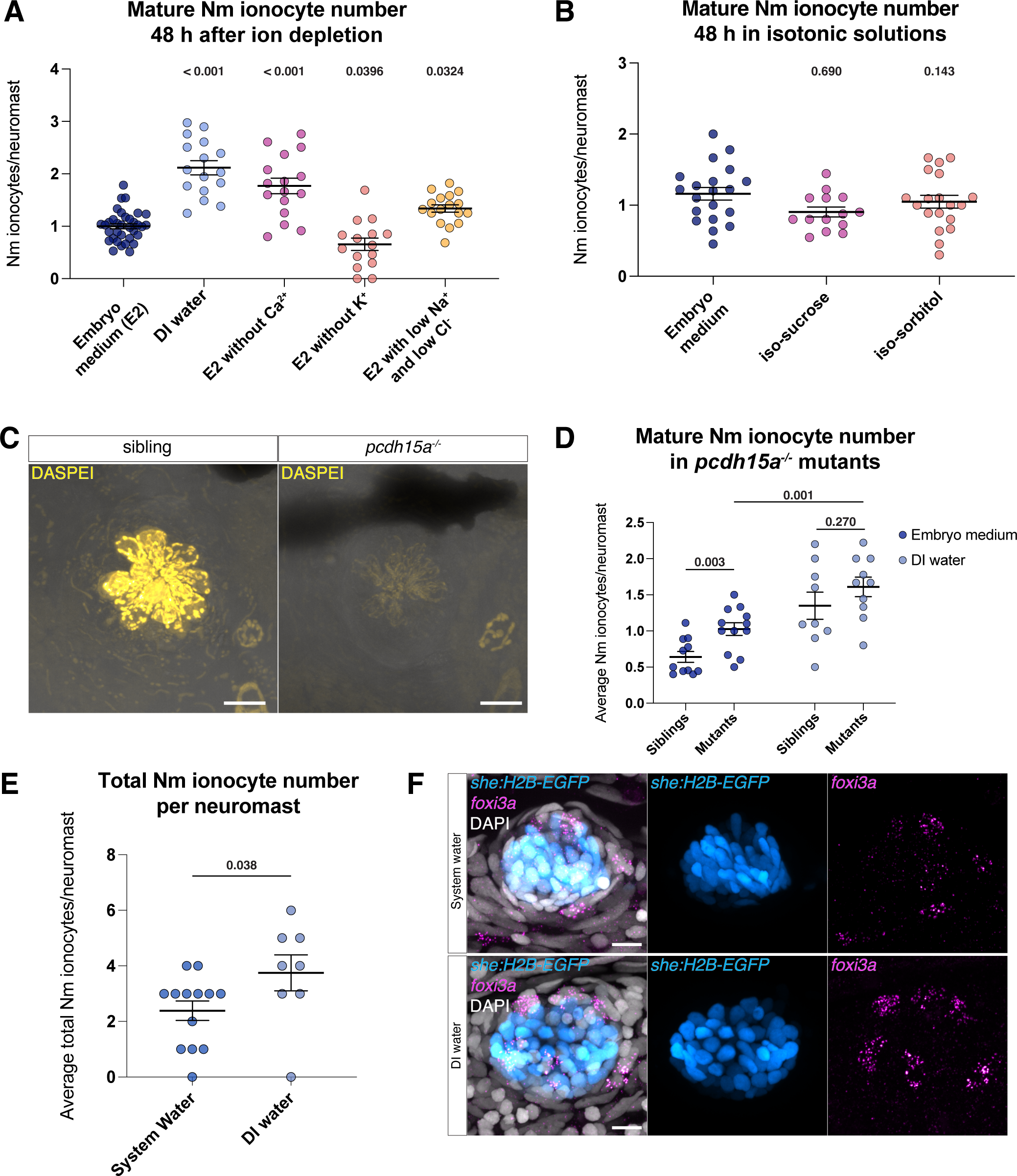
Nm ionocytes are triggered by low levels of Ca^2+^ and Na^+^/Cl^−^ ions. (A) Number of mature Nm ionocytes in regular embryo medium, DI water, and embryo medium depleted of either calcium, potassium or with low levels of sodium and chloride (embryo medium n= 34 larvae, DI water n = 16 larvae, no Ca^2+^ n = 16 larvae, no K^+^ n = 15 larvae, and low Na^+^/Cl^−^ n = 17 larvae, one-way ANOVA, normalized to embryo medium control). (B) Number of mature Nm ionocytes in isotonic solutions made with sorbitol or sucrose (embryo medium n= 20 larvae, DI water n = 18 larvae, iso-sorbitol n = 19 larvae, iso-sucrose n = 14 larvae, one-way ANOVA and Dunnett’s multiple comparisons test). (C) Active hair cells are labeled by DASPEI in *pcdh15a* siblings, but not in mutants. (D) Number of mature Nm ionocytes in siblings and mutants from *pcdh15a* mutants in embryo medium (Mann-Whitney test) and DI water. (E) Total Nm ionocytes number (progenitors and mature) in neuromasts from tailfin explants after 1h incubation in different salinities, (system water, n = 3 fish, 14 neuromasts, DI water, n = 2 fish, 8 neuromasts; Mann-Whitney test). (F) Representative images from (E).

Zebrafish, like other freshwater fish, live in a hypotonic environment and must balance the undesired uptake of water. Embryo medium (14-16 mOsmol/L) is hypotonic compared to zebrafish interstitial fluid (∼270 mOsmol/L), which causes undesired water influx that is compensated by the kidney^33^. To address if water influx is required for Nm ionocyte induction and differentiation, we incubated larvae in isotonic embryo medium supplemented with either D-sorbitol or sucrose to achieve interstitial fluid osmolarity^34,35^. Embryos raised in isotonic solution show similar numbers of Nm ionocytes compared to embryos raised in regular (hypotonic) embryo medium. This demonstrates that Nm ionocyte induction and differentiation does not depend on water influx caused by differences in osmolarity between the medium (hypotonic) and the fish (Figure 5B).

### Sensory hair cell activity controls Nm ionocyte number

We showed that the sensor is ion selective and osmolarity independent. We next asked which cells contribute to Nm ionocyte recruitment. Because Nm ionocytes modulate hair cell function and loss of ionocytes leads to a decrease in hair cell activity^18,36^, we asked if hair cells play a role in inducing more Nm ionocytes. Analysis of *pcdh15a* mutants, which develop non-functional hair cells (Figure 5C)^37^, shows that mutants have an increased number of mature Nm ionocytes irrespective of whether they were raised in embryo medium or DI water (Figure 5D). This suggests that disruption in hair cell activity plays a role in controlling number or recruitment of new Nm ionocytes. Interestingly, while inhibiting hair cell function recruits more Nm ionocytes, we see a further increase in Nm ionocyte number in mutants incubated in DI water compared to those incubated in embryo medium. This suggests that, even though hair cell function modulates the number of ionocytes recruited into neuromasts, there must be additional triggers of Nm ionocyte recruitment and differentiation.

### Salinity sensing and Nm ionocyte differentiation do not depend on systemic factors

Our *pcdh15* mutant analyses suggest that, in addition to hair cells, other cell types are involved in regulating Nm ionocyte number. Sensory neurons that innervate the skin and sense other environmental changes such as temperature and oxygen in other organisms could also be involved in this process^38–40^. *adgra2* mutants lack dorsal root ganglia (DRG) and somatosensory neurons^41–43^ but show no defects in progenitor activation or Nm ionocyte differentiation based on *foxi3a* and *trpv6* HCRs (Figure S7A and S7B). Therefore, we conclude that DRG-derived sensory neurons are not involved in sensing salinity and inducing ionocytes.

Another possibility is that basal cells or Nm ionocyte progenitors could autonomously sense changes in ionic concentration in the medium, as indicated by the fast upregulation *foxi3a* and *foxi3b* in basal cells upon salinity decrease. To test this hypothesis, we assayed Nm ionocyte number in neuromast-containing tailfin explants exposed to different salinities. Nm ionocyte numbers increased in explants incubated in DI water compared to those in system water (Figure 5E and 5F). These results demonstrate that either basal stem cell-derived progenitors or neighboring cells may directly detect salinity changes and that salinity sensing does not require sensory neurons or systemic signals, such as hormones. Overall, we conclude that Nm ionocyte number is controlled by multiple cell types and through distinct mechanisms.

## DISCUSSION

Physiological adaptation happens in individuals that are subjected to environmental perturbations during their lifetime. These changes are rapid and reversible, in contrast to natural selection-driven genetic adaption. One example is the adaptive differentiation of ionocyte subtypes within the body in response to fluctuations in water salinity, including Nm ionocytes^18^. In this study, we show that in response to a decrease in medium ion concentration, Nm ionocyte precursors, derived from basal stem cells of the skin, upregulate the transcription factors *foxi3a* and *foxi3b* and divide to give rise to a Nm ionocyte pair. Here, we discuss the possible sources of Nm ionocyte progenitors and their similarities and differences with other tissue-specific ionocytes. We also discuss possible mechanisms that sense environmental salinity and how comparisons between tissue-specific ionocytes may inform us about the evolution of cell types.

### Are Nm ionocyte progenitors a defined subpopulation of cells or are all basal cells competent to respond to environmental changes?

Basal cells are multipotent stem cells that give rise to several cell types in organs such as the lungs, skin and tastebuds^44–46^. Basal stem cells are heterogeneous populations and different subtypes or states have different potencies. In tastebuds, only a subpopulation of SOX2^high^ basal cells can differentiate into taste-receptor cells^47^ and in the pulmonary epithelium, subtypes of basal stem cell populations give rise to different lineages^48^. In the frog epidermis, basal cells differentiate into multiciliated cells, small secretory cells and ionocytes^49,50^, while in zebrafish basal cells give rise not only to skin and Nm ionocytes, but also to other cell types such as keratinocytes, mucous cells and Merkel cells^51–53^. The basal cell heterogeneity in other systems, combined with our observation that only a small population of cells rapidly respond to low salinity, leads us to hypothesize that Nm ionocytes are derived from a small subpopulation of primed basal stem cells.

Adult stem cells are maintained through asymmetric cell divisions to prevent the depletion of the stem cell pool throughout life^54–57^. We propose that a similar mechanism may act in the zebrafish skin to maintain pre-determined Nm ionocyte precursors (Figure S8, top right).

### Different mechanisms for sensing salinity

Low salinity leads to the adaptive differentiation of new Nm ionocytes. Our tissue explant experiments show that tissue-autonomous sensing is sufficient to induce Nm ionocyte differentiation. One possibility is that the local components that induce the increase of Nm ionocyte number in explants are hair cells. Another is that basal cells themselves sense environmental changes and respond accordingly. In the mammalian lung, TP63-positive airway stem cells sense hypoxia cell-autonomously and differentiate into solitary neuroendocrine cells, which mitigate hypoxic injury^58^. If basal cells are directly sensing ionic concentration, they likely do so through ion channels. However, it is still not known whether or which ion channels are present in Nm ionocyte progenitors, as we have not detected in progenitors the expression of the ion channels present in mature ionocytes.

Nevertheless, there is a possibility that systemic responses involving the olfactory system, nerves or hormonal regulation also play a role in salinity sensing. Zebrafish larvae sense NaCl concentrations via the olfactory epithelium, which impacts swimming behavior, but it remains unknown if the same circuit controls ionocyte number^59^. However, in contrast to what we observed in Nm ionocytes, the olfactory system only detects changes in salinity, not absolute levels, suggesting two distinct mechanisms. Sensory neurons in *Drosophila* detect environmental changes such as oxygen availability and in response regulate cell fate^39,40^. Here, we demonstrate that DRG-derived sensory neurons are not required for the response to low salinity as Nm ionocyte development is normal in *adgra2* mutants. However, Rohon-Beard neurons are still present in *adgra2* mutants and could potentially be involved in sensing environmental changes.

Disruption of hair cell function leads to an increase in Nm ionocyte number, indicating that hair cell activity is an important regulator of this process. However, because the number of Nm ionocytes increases even further when mutant larvae are incubated in DI water, additional mechanisms or cell types are likely at play. It has recently been suggested that lateral line hair cells do not rely on K^+^ influx for mechanotransduction but rather on an outward flow of anions instead^60^. In this case, Nm ionocytes could likely control the ionic microenvironment at the basolateral cell surfaces of hair cells. One hypothesis is that hair cells directly influence basal cell differentiation based on ionic availability in their microenvironment. The mechanistic relationship between Nm ionocyte number and hair cell activity, and how hair cell function may feedback on Nm ionocyte numbers will be focus of future work.

### Ionocyte number is controlled by different mechanisms in the skin and neuromast

We showed that Ca^2+^ depletion leads to formation of new Nm ionocytes through activation of new progenitors. Likewise, lowering the concentrations of Na^+^ and Cl^−^ in the medium leads to an increase in Nm ionocyte number, but to a lesser extent than Ca^2+^ depletion. On the other hand, K^+^ depletion is not sufficient to induce new Nm ionocytes. These results suggest that not all ions play the same role in Nm ionocyte recruitment. Interestingly, the increase of Nm ionocytes in each condition did not correlate with the ion concentration in normal embryo medium: the concentrations of Na^+^ and Cl^−^ are more than ten-fold those of Ca^2+^ and K^+^, but Ca^2+^ removal leads to the strongest response.

Ca^2+^ depletion induces ionocyte proliferation in both skin and neuromasts. In neuromasts, low Ca^2+^ leads to formation of a new pair of Nm ionocytes through division of a progenitor cell, which contrasts with the skin where most NaR ionocytes are solitary. Skin NaR ionocytes remain quiescent during homeostasis due to intracellular Ca^2+^-dependent suppression of PI3K, and this intracellular Ca^2+^ is imported to the cell via the Trpv6 channel^61^. Under low Ca^2+^ conditions, NaR ionocytes exit quiescence and proliferate^61–64^. Although Nm ionocyte progenitors also proliferate under low Ca^2+^, they do not express *trpv6*. Mature Nm ionocytes, on the other hand, express *trpv6* but are post-mitotic. These observations suggest that while low Ca^2+^ induces more ionocytes in both skin and neuromasts, it does so through different mechanisms.

### Notch signaling dynamics differ between skin and Nm ionocyte development

Notch signaling governs the specification and differentiation of several types of ionocyte in different organisms, from fish and frogs to mice and humans^20,21,50,65^. In the frog epidermis, Notch inhibition leads to an increase in ionocyte number, similar to the zebrafish embryonic skin^66^. In the kidneys of mammals, Notch signaling also specifies ion-regulating cells^67,68^. We show that the Nm ionocyte pair is comprised of a cell with low Notch (*dll4*-positive) and a cell with high Notch signaling (*notch1b-*positive), and that Notch signaling is sustained in mature, fully differentiated ionocytes (Figure 2)^18^. This contrasts with skin ionocyte differentiation in fishes and frogs, where Notch signaling is expressed at the onset of fate determination and is not maintained in mature cells. In the embryonic zebrafish skin, Notch-dependent lateral inhibition leads some basal stem cells to express *dlc* and *jagged* ligands, surrounded by cells that express *notch1a* and *notch3* receptors^20,21^. This results in the differentiation of the low Notch cell (*dlc*-positive) into a skin ionocyte, while the Notch-positive cell becomes a keratinocyte (Figure S8, left panel).

The different roles played by Notch signaling in skin and Nm ionocyte development are apparent in loss- and gain-of-function experiments. Global downregulation of Notch signaling in *mindbomb* mutants or pharmacological Notch inhibition lead to an increase in skin ionocyte density at expense of keratinocytes^20,69^. In contrast, pharmacological inhibition or loss of Notch signaling in *dll4* mutants causes loss of mature Nm ionocytes, but no changes in the progenitor population (Figure 3)^18^. These results demonstrate that while Notch signaling is crucial for proper ionocyte specification, differentiation or survival, the role played by Notch are context- and tissue-dependent.

While *dll4* plays a pivotal role in Nm ionocyte survival, it had not been shown to have a role in skin ionocytes. Here, we show that *dll4* mutants lack the NaR ionocyte population in the skin, concomitant with an expansion of the HR population. These two subtypes of ionocytes share a common ionocyte progenitor (Figure S8), and little is known about how this progenitor decides between the NaR or HR fate. It has been suggested that *foxi3b* levels drive differentiation of each subtype, but this has not been demonstrated^20^. Our results suggest that *dll4* may play a role in this decision, and its loss shifts the population entirely to the HR fate. An alternative possibility is that, similar to Nm ionocytes, NaR ionocytes also undergo cell death in the mutants, and the HR population increases to compensate for their loss.

Ion regulating cells exist in many invertebrate and vertebrate species^4,11,12,50,70–73^. While homology between these cell types has not been established, mammalian ionocytes express the *foxi3a/b* ortholog Foxi1. Likewise, hemichordates (acorn worms) and cephalochordates (amphioxi) possess foxi1-positive cells in their gills, which are hypothesized to be proto-ionocytes^74^. These putative ionocytes express several of the ion channels present in zebrafish ionocytes. Therefore, appearance of bona-fide ionocytes likely predates the evolution of vertebrates and of a lateral line system. The emergence of tissue-specific ionocytes in vertebrates serves as an excellent new model to understand the evolution of cell type diversification^75^.

### Limitations of this study

We showed that Nm ionocyte differentiation is associated with the upregulation of Foxi3a/b transcription factors and a temporally regulated expression of Notch pathway components. However, most of our analyses relied on the analysis of fixed samples, and the temporal resolution of genes may be more dynamic than what we can assess. Additionally, while it is still likely that Notch disruption leads to a fate change in the Nm ionocyte pair, we do not have markers to assess fate before cells die. We can therefore only conclude that loss of Notch signaling affects cell survival, but it is possible that Notch signaling also plays a role in fate determination.

## ACKNOWLEDGMENTS

We are grateful to Paloma Meneses-Giles, Daniela Münch, Drs. Alice Accorsi, Aurelie Hintermann and Cameron Berry for critical reading of the manuscript and all members of the Piotrowski lab for insightful discussions. We also thank Dr. J. Rasmussen and T. Nicolson for fish lines, Shin-Ichi Higashijima for the Gbait plasmid, Sean McKinney for his help with image analysis, the Stowers Institute Aquatics Facility for fish care, and the Stowers Media Prep facility, particularly Rebecca Stephenson, for providing embryo media with different modifications. The work was funded by a NIH (NIDCD) award 1R01DC015488-01A1 to T.P. and by institutional support from the Stowers Institute for Medical Research to T.P.

## AUTHOR CONTRIBITIONS

Conceptualization: JP and TP; Methodology: JP and TP; Investigation: JP, MEL, YT; Data Analysis: JP, CW; Writing – original draft: JP and TP.; Writing – review and editing: JP, MEL, YT, CW, TP; Funding acquisition: TP; Resources: TP; Supervision: TP

## COMPETING INTERESTS

No competing interests declared.

## RESOURCE AVAILABILITY

### Lead contact

Further information and requests for resources and reagents should be directed to and will be fulfilled by the Lead Contact, Tatjana Piotrowski.

### Materials availability

Fish lines generated in this study are available from the lead contact.

### Data and code availability

Original data underlying this manuscript can be accessed from the Stowers Original Data Repository at http://www.stowers.org/research/publications/libpb-2446

## MATERIALS AND METHODS

### Zebrafish lines and husbandry

All experiments were performed following the guidelines of the Stowers Institute IACUC review board. Embryos were raised in 0.5x embryo E2 medium (7.5 mM NaCl, 0.25 mM KCl, 0.5 mM MgSO_4_, 75 mM KH_2_PO4, 25 mM Na_2_HPO_4_, 0.5 mM CaCl_2_, 0.5 mg/L NaHCO_3_, pH = 7.4) unless specified otherwise. Wild-type fish used were either AB or TU. The following published transgenic lines were used in this study: *Tg(−8.0cldnb:lynEGFP)^zf106Tg^* ^76^, *Tg(−8.0cldnb:H2A-mCherry)^psi4Tg^* ^77^, *Tg(tp1bglobin:EGFP)^um13^* ^78^, *Tg(she:H2B-EGFP)^psi59Tg^* ^18^.

### Mutant line generation

Mutants for *dld* and *dll4* genes were generated using CRISPR/Cas12a. For the *dll4^psi^*^87^ mutation, two gRNAs targeting sequences 869 bp upstream of the translation start site (5’-GTATCCCAGAAAACACTATA-3’) and in Exon 4 (5’-GTTGCAGATGAACCGGTAAG-3’) were designed using DeepCpf1^79^ and checked for off-target effects using CRISPRscan^80^. For the *dld^psi^*^86^ mutant, two CRISPR/Cas12a guides were designed using the same parameters and tools, and targeted regions 382 bp upstream of the translation start site (5’-GATCATCTCCACATGAACTT-3’) and 215 bp downstream of the stop codon (5’-CCAGGAACGTGCCAAATGGA-3’).

For both mutants, one cell stage zebrafish embryos were injected with 50 nM EnGen® Lba Cas12a (Cpf1) (New England Biolabs) and each gRNA at a final concentration of 2 nM. Injected larvae were raised to adulthood and germline transmission of the mutation was confirmed by PCR using the following primer pairs: *dll4*-FwA: 5’-TTTGGGATTCACCTTGAACG-3’, *dll4*-FwB: 5’-GCCTGGCACTCACCTTACTC-3’, *dll4*-Rv: 5’-ATAACTGGCCATCTGGGTTG-3’, *dld*-Fw: 5’-GCCTGTATATAACCGGCTGC-3’, *dld*-RvA: 5’-GTCAGTGCATATGCATACACAG-3’ and *dld*-RvB: 5’-TCATTAGTCGTCCCATGGCG-3’.

Once F0 founders were identified, one founder per gene was outcrossed to wild-type fish to generate a stable F1 generation. The F1 generations were then raised to adulthood and heterozygous carriers of the deletion were identified by genotyping PCR from fin clips. Experiments were performed with embryos derived from incrosses of either F1 or F2 parents.

### Transgenic line generation

The transgenic line *Tg(hsp70l:dld;cmcl2:EGFP)^psi88Tg^* was generated using the tol2 kit^81^. The cDNA sequence for *dld* was cloned from isolated cDNA using Phusion High Fidelity Polymerase (New England Biolabs) and cloned into a pME vector digested with EcoRI using Gibson Assembly and the following primers for homology: 5’-GTATCGATAAGCTTGATATCGAATTCACCATGGGACGACTAATGATAGCTG-3’ and 5’-GTGGATCCCCCGGGCTGCAGGAATTTTACACCTCGGTTGCAATAATAC-3’.

*Tg(dld:hist2h2l-EGFP)^psi84Tg^* knock in was generated via CRISPR-knock in based on a previously published method^82^ with modifications for Cas12a. The Gbait vector was first modified via replacement of the Gal4FF-BGH-polyA-Kanamycin resistance sequence (BamHI/SalI digested) with a *hist2h2l-EGFP-sv40* polyA fragment (BglII/XhoI digested).

A Cas12a targeting sequence, 5’-ACACAGGAAACAGCTATGAC-3’, was identified within the GBait plasmid upstream of the T3 primer sequence. This sequence is not predicted to target the zebrafish genome. We verified that this guide RNA digested the Gbait plasmid in combination with Lba Cas12a (New England Biolabs) in an *in vitro* reaction (data not shown). To identify gene specific guides, DeepCpf1^79^ was used, followed by BLAST of the zebrafish genome to ensure specificity. A guide (5’-GAGATGAAAACTTCAAACT-3’) was identified that would target 940bp upstream of the 5’UTR of *dld*. All guide RNAs were ordered from IDT, then diluted to a stock concentration of 24uM^83^.

To make the injection mix, the Gbait and *dld* gRNAs were combined at 4.8uM each with 20uM Lba Cas12a protein (New England Biolabs) and incubated for 10 minutes at 37^0^C. The mix was then placed on ice and 30ng of Gbait-H2B-GFP plasmid and phenol red were added. We injected 1-3nl of the mix into the cell of one-cell stage zebrafish embryos. Embryos were raised at 28°C then screened for GFP expression at 1-2dpf. GFP positive larvae were raised to adulthood then screened for germline transmission of GFP.

## METHOD DETAILS

### Hybridization Chain Reaction

Hybridization chain reaction (HCR) was adapted from the manufacturer’s instructions^84^ (Molecular Instruments). Embryos were fixed overnight with 4% PFA at 4°C, then dehydrated in an ethanol series up to 100% ethanol. Dehydrated samples were stored at −20°C from overnight up to 6 months. Samples were then rehydrated and washed twice with H_2_O+0.1% Tween20 (Sigma-Aldrich) and incubated in 80% acetone at −20°C for 20 minutes. The samples were then washed twice in room temperature (RT) PBST (PBS + 0.1% Tween20) and incubated in 200uL of Hybridization Buffer (Molecular Instruments) for 30 minutes at 37°C. Later, samples were incubated in Hybridization Buffer with 2pmol of each probe at 37°C overnight. On the next day, samples were washed 4 times for 15 minutes with pre-warmed Washing Buffer (Molecular Instruments) and twice in 5x SSCT for 5 minutes, and subsequently incubated for at least 30 minutes with 200uL of Amplification Buffer equilibrated at RT. Finally, samples were incubated overnight with the suitable amplifiers for each experiment, then washed 3 times with 5x SSCT and incubated with DAPI (1ug/mL). Samples were protected from light and at 4°C prior to imaging. The following probes were used in this study: *notch1a*-B2*, notch1b*-B2*, notch3*-B2*, dla*-B1*, dlb*-B1*, dlc*-B1*, dld*-B5*, dll4*-B1*, jag1a*-B1*, jag1b*-B1*, jag2a*-B1*, jag2b*-B1, *her15*-B2, *hes6*-B1, *her4*-B2, *hey1*-B4, *lfng*-B2*, foxi3a*-B2*, foxi3b*-B4*, trpv6*-B1*, fosab*-B4*, junba*-B1, *dusp1*-B2, *kcnj1a*-B2, *rhcgb*-B5, *slc4a1b*-B1. The *her4*, *kcnj1a* and *her15* probes do not distinguish between tandem duplicates due to sequence similarity. Amplifiers were conjugated either with Alexa488, Alexa546 or Alexa647.

### Immunohistochemistry

Zebrafish larvae at different stages were anesthetized with MS222 (160 mg/mL) and fixed overnight at 4°C with 4% PFA. Standard antibody staining was performed for a5 (DSHB, AB_2166869)^85^.

### Cell Proliferation analysis

Larvae were treated for 24h with 3.3 mM EdU (Carbosynth) with 1% DMSO in E2, then fixed in 4% paraformaldehyde overnight at 4°C. Larvae were then stained with HCR and EdU simultaneously^86^. Briefly, HCR protocol was performed as described above, until the probe wash step. Then, EdU was clicked with 2mM CuSO4, 0.4% Triton-X, 2.5 uM 594-Azide and 50 mM ascorbic acid for 30 minutes and washed three times for 10 minutes with PBT (PBS + 0.8% Triton-X). After the EdU developing reaction, samples were incubated in HCR amplification buffer (Molecular Instruments) for 30 minutes and incubated overnight at room temperature with B4-647 amplifier.

### Acclimation of zebrafish larvae to different salinities and media

For acclimation in different media, fish were raised in embryo medium at a density of 40 embryos per dish until 3 dpf, with daily media changes. After reaching 3 dpf, fish were incubated in fresh, regular or modified embryo medium. For calcium depletion, embryo medium was made without CaCl_2_; for potassium depletion, embryo medium was made without KCl and KH_2_PO_4_; for sodium and chloride depletion, no NaCl, Na_2_HPO_4_ or NaHCO_3_ were added to the media. Isotonic solutions (270mOsmol/L) were made with either 242mM sorbitol or 238mM sucrose. Media pH was always adjusted to 7.2 on the same day of usage. Fish were incubated in different media for 48h at 28.5°C unless specified otherwise.

### Quantification of Nm ionocytes

Mature Nm ionocytes were identified and counted with double HCRs for *foxi3a* and *trpv6*, while progenitors were considered as *foxi3a*-positive, *trpv6-*negative cells adjacent to neuromasts and with a crescent morphology. Neuromasts from the anterior lateral line (Ml1, O2, Ml2, IO4, O1) and posterior lateral line (L1-L3, LII.1, LII.2) were quantified^6,87^. Numbers were averaged per larvae and displayed as one value that could also be lower than 1 if not all neuromasts contained ionocytes. For ionocyte frequency quantifications, neuromasts were classified based on the presence and absence of ionocytes and data is displayed as percentage of neuromasts that contained ionocytes. In ion depletion experiments, data between replicates have a wide distribution, and comparison between replicates was challenging. In this case, data was normalized based on each replicate’s embryo medium control. Normalized data are indicated in the respective figure legend.

For Nm ionocyte progenitors, both individual and pairs of *foxi3a/b-*expressing cells were counted as one progenitor unit. Cells were also only considered progenitors when they were immediately adjacent to neuromasts and not expressing the ion channel *trpv6*. When pairs were already expressing *foxi3a/trpv6* and were fixed while invading neuromasts, they were considered as mature ionocytes.

### Time-lapse and confocal imaging

Images were acquired on a Nikon Ti2 Yokogawa CSU-W1 spinning disk head equipped with a Hamamatsu Orca Fusion sCMOS. Objective lenses used were CFI Apo LWD 40x WI 1.15 NA Lambda S and CFI Apo 20x WI 0.95 NA Lambda S.

For live imaging experiments, larvae were slowly anesthetized with tricaine (MS222) up to 150mg/L and mounted in glass bottom dishes (Cellvis) using 0.8% low melting point agarose dissolved in either embryo medium or system water supplemented with MS222 (120mg/L)^88^. A Stage Top Incubator (OkoLab) was used to keep the constant temperature of 28.5°C and 85% humidity.

A Nikon LUNV solid state laser launch was used for lasers 395/405, 488, 561 and 647nm for GFP/Alexa488, RFP/Alexa546, and Alexa647 respectively. Emission filters used on the Nikon were 480/30, 535/30, 605/52. Nikon Elements Advanced Research v5.41.02 (Nikon) was used for image acquisition.

### Statistical analysis

All statistical tests and plots were performed in GraphPad Prism 10 (version 10.0.2). *p-* values smaller than 0.05 were considered significant. Normal distribution was assessed using Kolmogorov-Smirnov test. Data from two groups were compared using two-tailed unpaired t-test or two-tailed Mann–Whitney U test. When comparing data from more than two groups, statistical significance was calculated using either one-way ANOVA with Dunn’ s multiple comparison test or non-parametric Kruskal-Wallis ANOVA with Dunn’s multiple comparison test. Plots were made in GraphPad Prism 10 or the R package ggplot2^89^.

### Image analysis

Fluorescence intensity quantification of transgenic lines and HCRs (*dusp1*, *fosab* and *junba* HCRs) were performed in Fiji^90^. Images were Z-projected (maximum intensity projection), background was removed with the “Subtract Background” function, and a rolling ball radius of 50 pixels was used for all images and measured channels. Then, a ROI was drawn around the cell or cells of interest and measured using the “Measure” function. Values were then exported and the mean fluorescence intensity per ROI was imported into GraphPad Prism 10 for statistical analysis and graphing.

For HCR quantification of Notch receptors and putative downstream factors, we automatically segmented Notch-positive Nm ionocytes using the *Tg(tp1bglobin:EGFP)^um13^* transgenic line. To segment the ionocytes, the image file was opened with the python package tifffile^91^ and the ionocyte channel was preprocessed with background subtractions and smoothing. A 3D white tophat (scipy)^92^ with a footprint of size of 50×50×3 pixels was used for background subtraction, and a 3D gaussian filter with a sigma of 1×1×0.5 was used for smoothing. After preprocessing, a threshold was applied to the image at a grey value of 500 to produce a segmentation mask. To slightly reduce the width of each mask, an Euclidian distance transform was calculated and pixels with transform values less than 2 were set to zero, while pixels with value 2 or greater were set to one. The resulting mask will often have multiple parts that merge into one structure at the bottom of the z-stack, since the Notch reporter also labels central support cells in the neuromast. To divide and segment individual cells, the z-stack was treated as a timelapse, and each cell was tracked in z. Individual cells were determined by only using trajectories before merge events. With the desired cells segmented, the other channels were background subtracted by white tophat with a footprint of (15, 15, 1) and the mean intensity within each cell was measured.

Quantification of fluorescence in either the *dld:H2B-EGFP* or asymmetry in *dld* expression via HCR was done in Fiji. In these cases, either the brightest slice or a maximum intensity projection of up to three brightest slices were quantified. Background was not subtracted but the comparison was performed pairwise with the cell from the same image.

In the case of some reporter lines such as the *dld:H2B-EGFP* line, fluorescence intensity is heterogeneous between cells. Therefore, we applied nonlinear changes (gamma) to make all features visible. Gamma ranged from 0.5 to 0.8 and was applied whenever stated in figure legends. Image quantification was always performed in the raw file and never with gamma-modified images.

---

**Figure S1:**
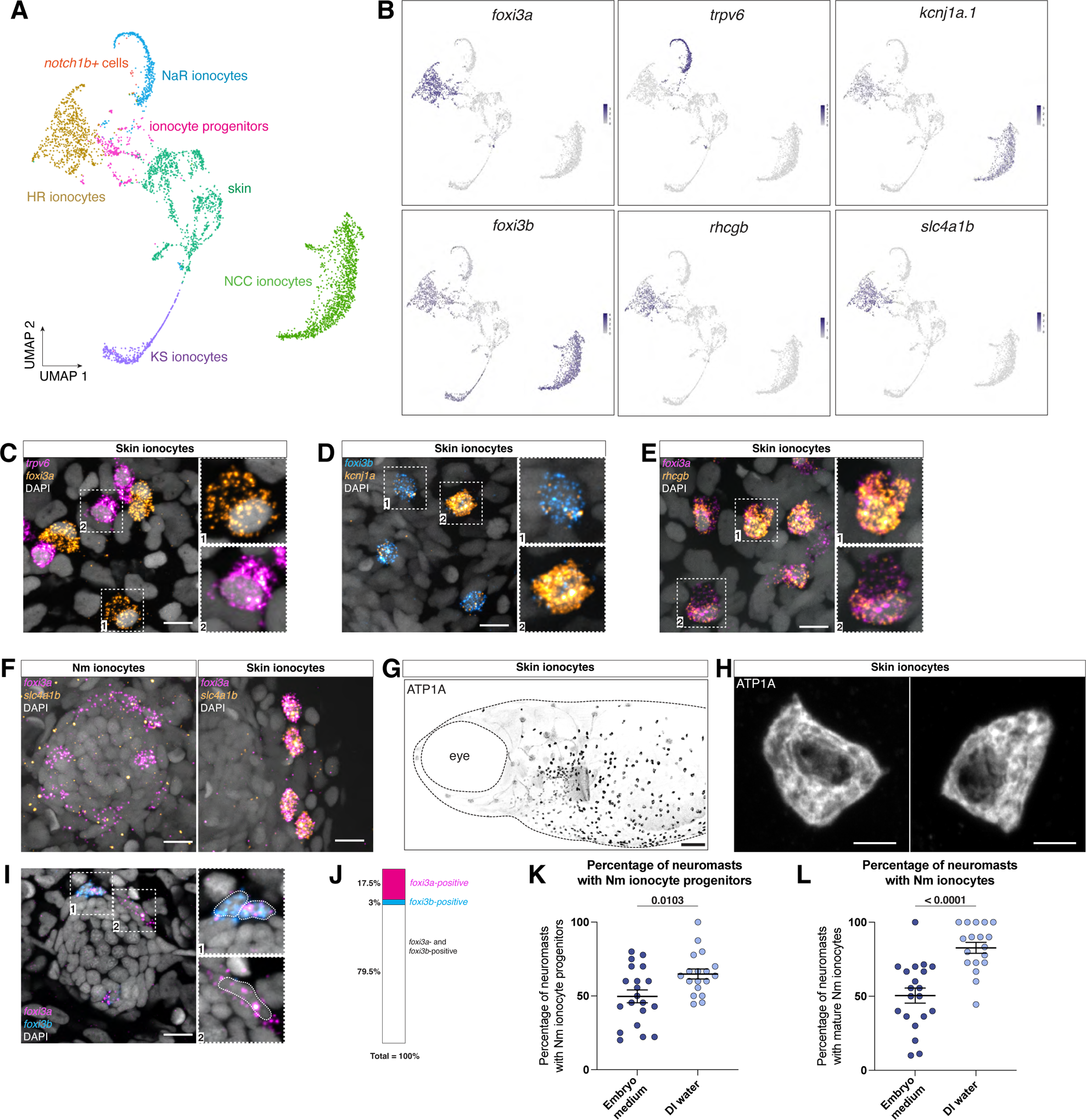
Differences in gene expression and morphology between skin and Nm ionocytes. (A) UMAP showing different clusters of ionocyte subtypes from Peloggia and Muench et al., 2021. (B) Feature plots for markers of different ionocyte subtypes (C) Maximum intensity projection of skin HR ionocytes, labeled by *foxi3a*, and NaR ionocytes, labeled by *trpv6.* (D) Some skin ionocytes, labeled by *foxi3b* HCR, express *kcnj1a* (E) Some skin ionocytes express *rhcgb*. (F) Nm ionocytes, mature or progenitors (left), do not express *slc4a1b*, but skin ionocytes do (right). (G) Whole mount 5 dpf zebrafish larva in which ionocytes are stained with Na(+) K(+) antibody. (H) High magnification images of (G) depicting skin ionocyte morphology. Scale bar = 5 µm. (I) Neuromast containing two sets of Nm ionocyte progenitors. In “1”, both *foxi3a* and *foxi3b* are expressed in both cells of the progenitor pair, while in “2” the progenitor is a single cell expressing only *foxi3a*. (J) Percentage of cells that co-express both transcription factors, *foxi3a* and *foxi3b*, and of cells that express only one out of the two factors. (K) Frequency of Nm ionocyte progenitors in different media depicted as the percentage of neuromasts in a larva that contain one or more Nm ionocyte progenitors (n = 18 larvae, 170 neuromasts; unpaired t-test). (L) Similar to analysis as in (K) for mature Nm ionocytes (unpaired t-test). Scale bars = 10 µm unless specified otherwise.

**Figure S2:**
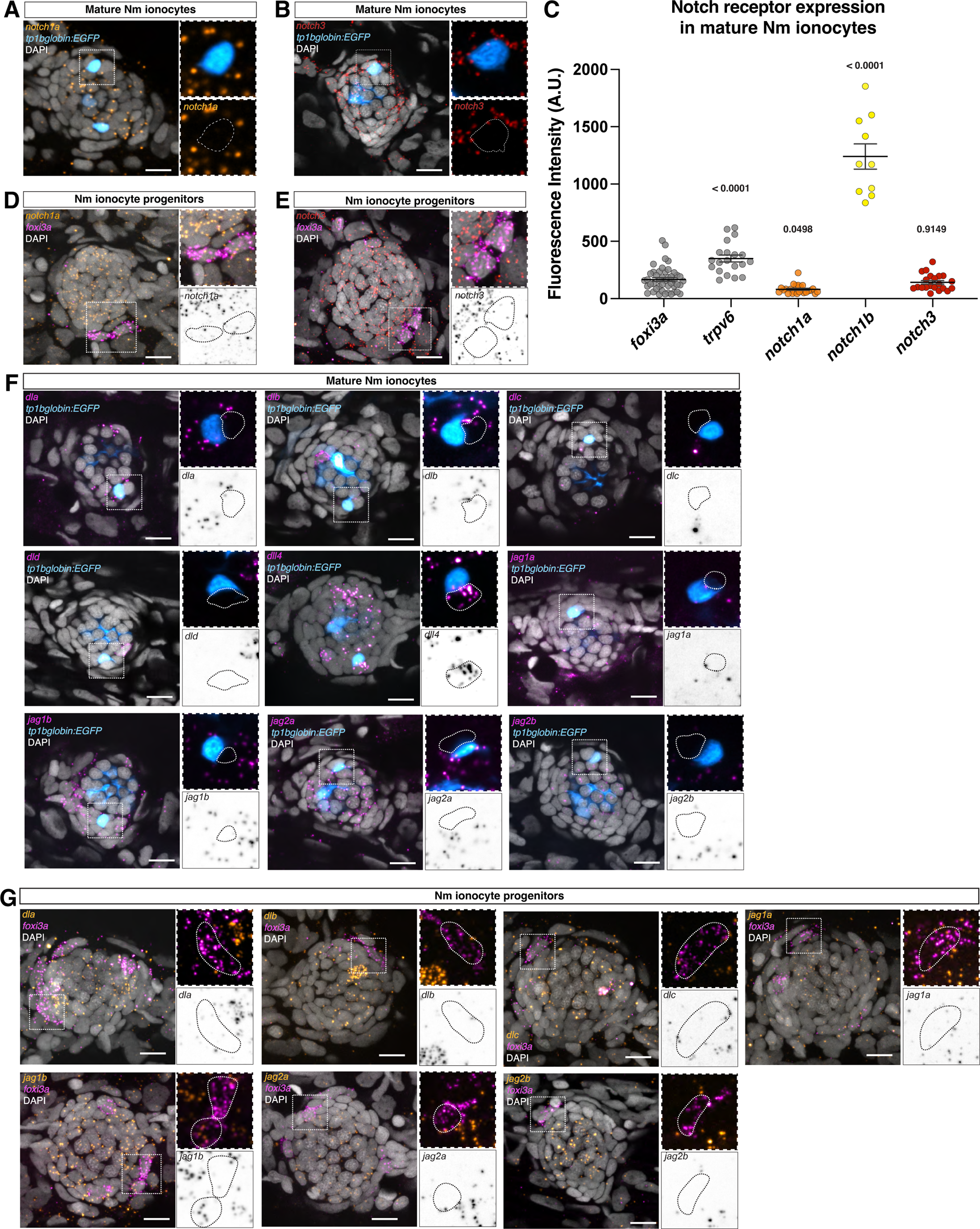
Notch ligand and receptor expression in Nm ionocytes. (A) HCRs for *notch1a* and (B) *notch3* in mature ionocytes. (C) Quantification of A, B and Figure 2A. *foxi3a* HCR serves as a negative control and *trpv6* HCR is used as a positive control of signal in Notch reporter-positive cells. Quantification was performed using the Notch reporter channel as a segmentation mask (see Methods) (D) Lack of expression of *notch1a* and (E) *notch3* in Nm ionocyte progenitors. (F) HCR for all other Notch ligands present in zebrafish in mature and (G) Nm ionocyte progenitors.

**Figure S3:**
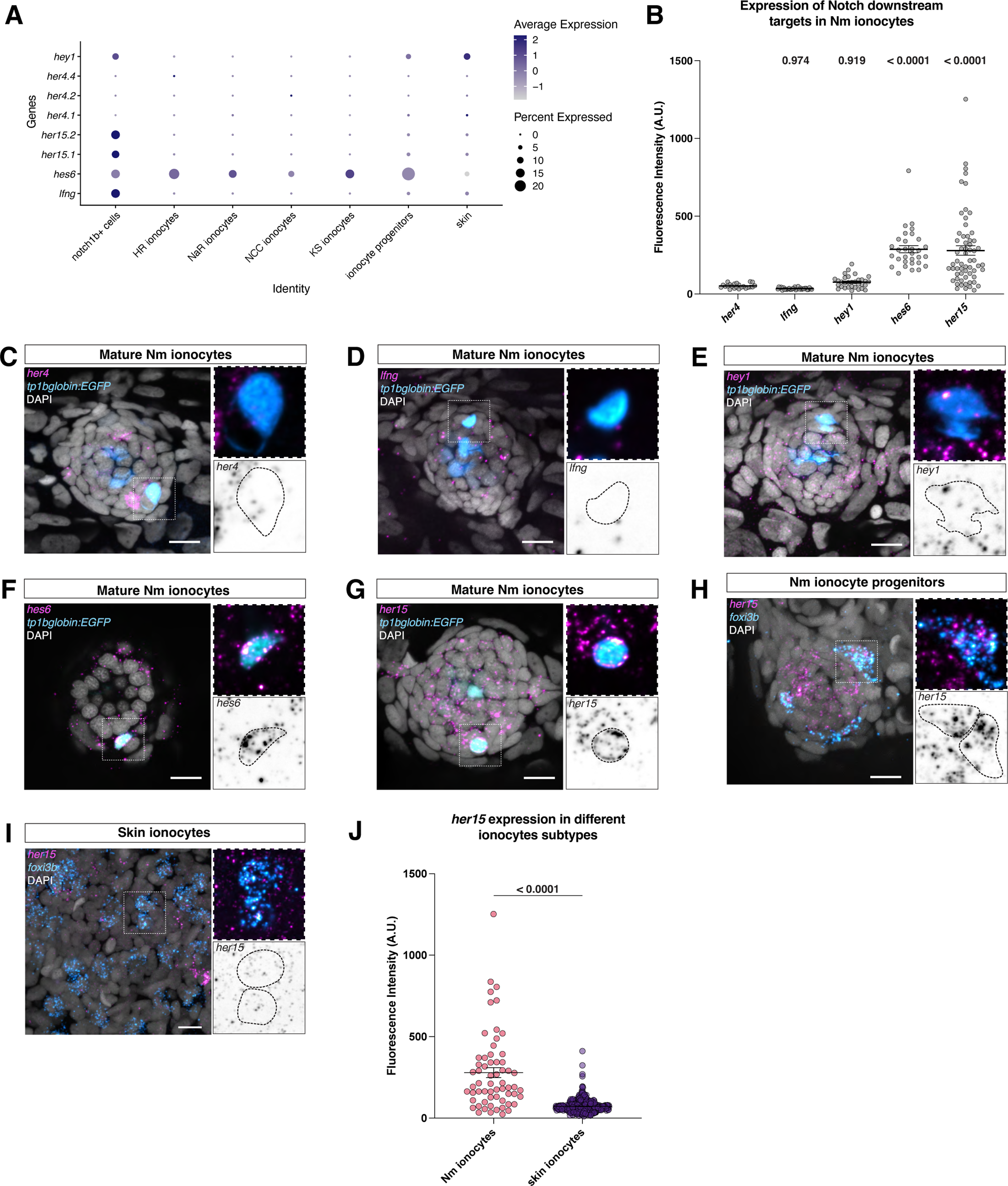
Analysis of Notch downstream target genes in Nm ionocytes reveals putative Nm ionocyte specific targets. (A) Dot plot from clusters on Figure S1A depicting candidate Notch downstream factors present in Nm ionocytes (*notch1b+* cells). (B) Quantification of panels (C-G), which show expression of Notch targets in mature Nm ionocytes. *her4* is used as a negative control, as it is not detected in the scRNA-seq data (C) HCRs for *her4*, (D) *lfng*, (E) *hey1*, (F) *hes6*, and (G) *her15*. (H) *her15* expression in Nm ionocyte progenitors and (I) in the skin. (J) Quantification of *her15* expression in skin ionocytes when compared to Nm ionocytes (Nm ionocyte data from (B)).

**Figure S4:**
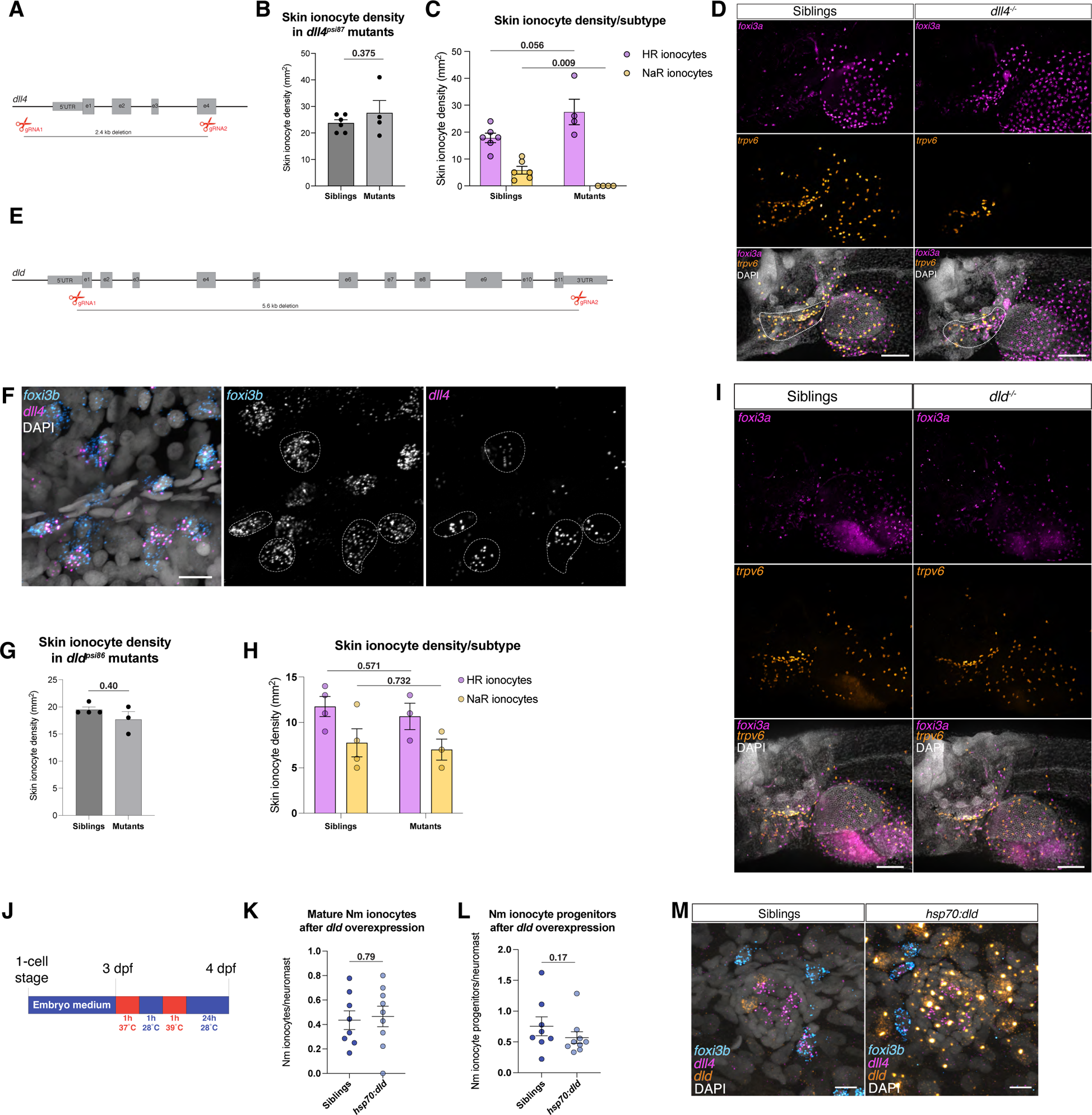
Manipulation of the Notch pathway shows a role for *dll4* in ionocyte development. (A) Schematic of *dll4* gene deletion strategy using CRISPR/Cas12a (B) Quantification of total skin ionocyte density in the yolk sac region of 5 dpf zebrafish larvae (Unpaired t-test) (C) Data from (B) split into HR and NaR ionocyte subtypes (Unpaired t-test). (D) Representative images from (C). Gills are outlined in dashed white. (E) Schematic of *dld* gene deletion strategy using CRISPR/Cas12a. (F) Quantification of total skin ionocyte density in the yolk sac region of 5 dpf zebrafish larvae between siblings and mutants (Unpaired t-test) (G) Data from (F) split into HR and NaR skin ionocytes. (H) Representative images from (G). (I) Heat shock paradigm used to upregulate *dld* in *Tg(hsp70:dld)* fish. (J) Number of mature Nm ionocytes (siblings, n = 8 larvae, *hsp70:dld*, 9 larvae, Unpaired t-test) and (K) progenitors 24h after heat shock (Mann-Whitney test, *p-value = 0.17*). (L) HCRs of fish fixed immediately after heat shock shows specific upregulation of *dld* gene in the transgenic. Scale bars = 10 µm.

**Figure S5:**
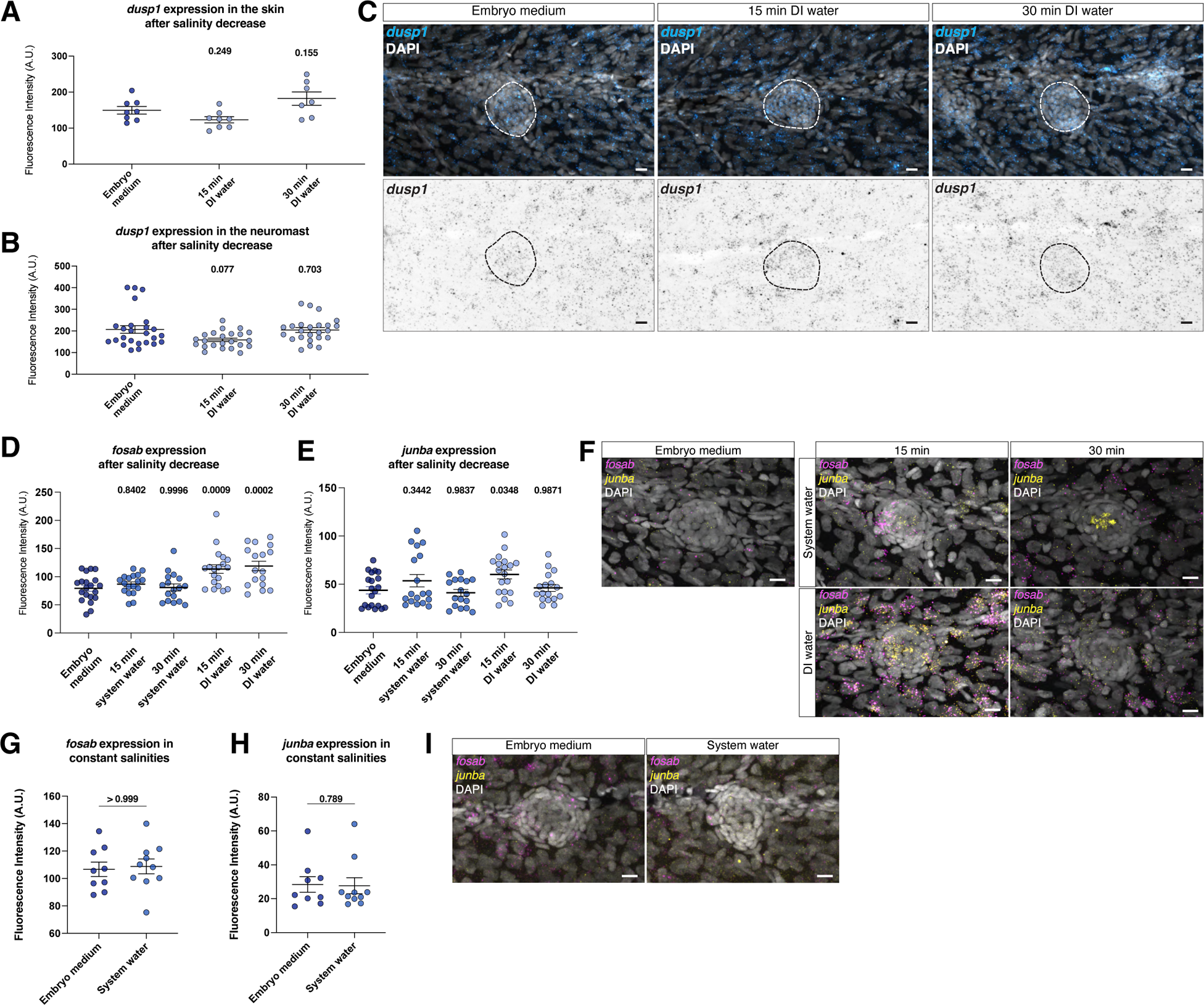
Nm ionocytes are not induced by stress response pathways. (A) Expression of the GR target gene *dusp1* in the skin and (B) neuromasts quantified by HCR at different times after low salinity incubation (embryo medium n = 4 larvae, 25 neuromasts, 15 min n = 5 larvae, 23 neuromasts and 30 min n = 5 larvae, 24 neuromasts, Kruskall-Wallis ANOVA) (C) Representative images from A and B. (D) Expression of *fosab* and (E) *junba* after 15 and 30 min of salinity decrease (Kruskall-Wallis ANOVA with Dunn’s multiple comparisons test). (F) Representative images from (C) and (D). (G) Expression of *fosab* and (H) *junba* in embryos raised in different salinities from the one cell stage (Mann-Whitney and Unpaired t-test, respectively). (I) Representative images from (G) and (H).

**Figure S6:**
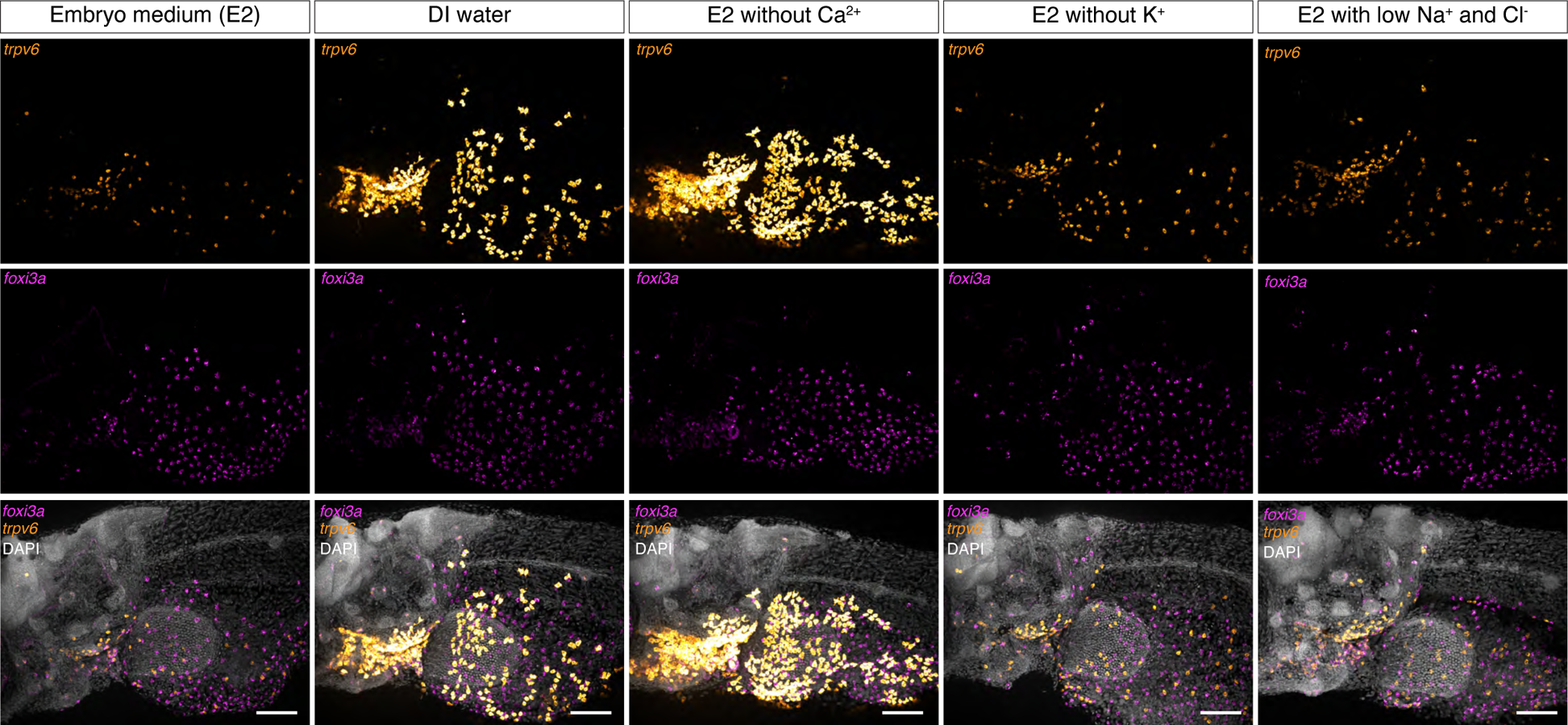
Skin ionocytes also respond to specific ion depletions. *foxi3a* (HR ionocytes) and *trpv6* (NaR ionocytes) HCRs showing response of HR and NaR ionocytes, to 48h incubation in embryo medium, DI water, depletion of Ca^2+^, K^+^ and low Na^+^/Cl^−^.

**Figure S7:**
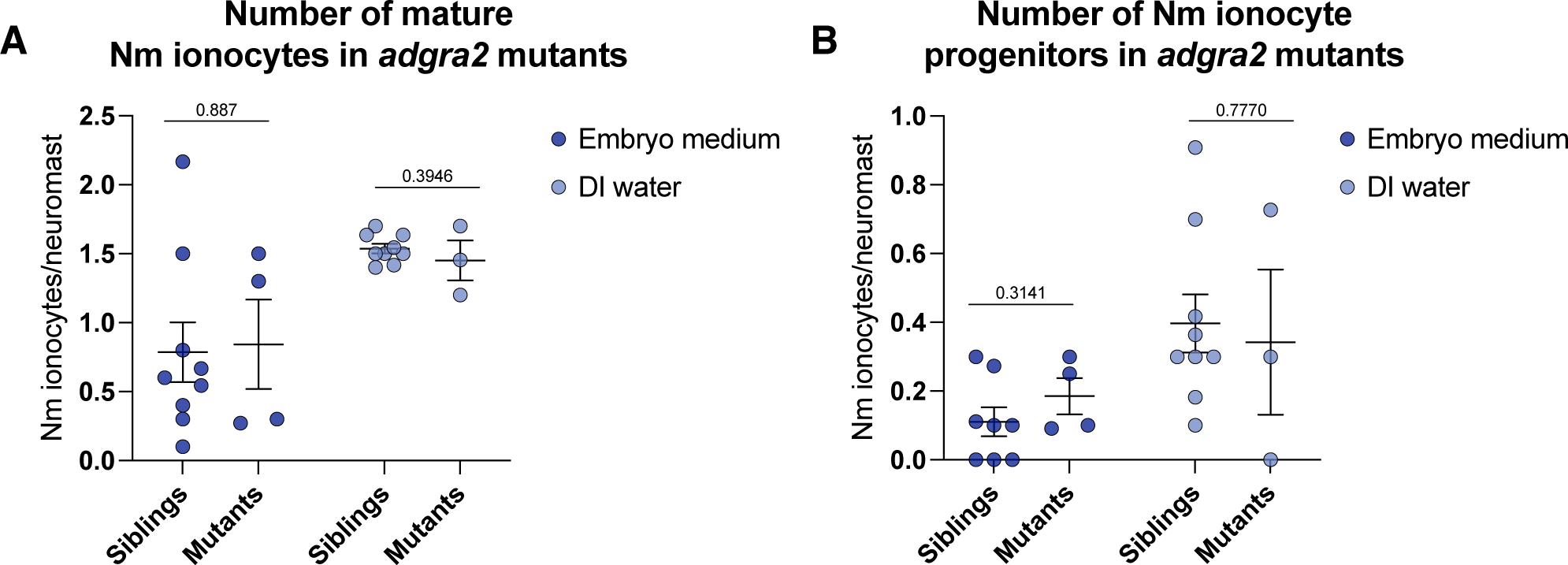
Nm ionocyte recruitment does not require sensory neurons. (A) Number of mature Nm ionocytes (B) and progenitors in *adgra2* mutants (siblings in embryo medium, n = 9 larvae, 86 neuromasts, mutants in embryo medium, n = 4 larvae, 43 neuromasts, siblings in DI water, n = 9 larvae, 94 neuromasts, mutants in DI water, n = 3 larvae, 31 neuromasts, unpaired t-test).

**Figure S8:**
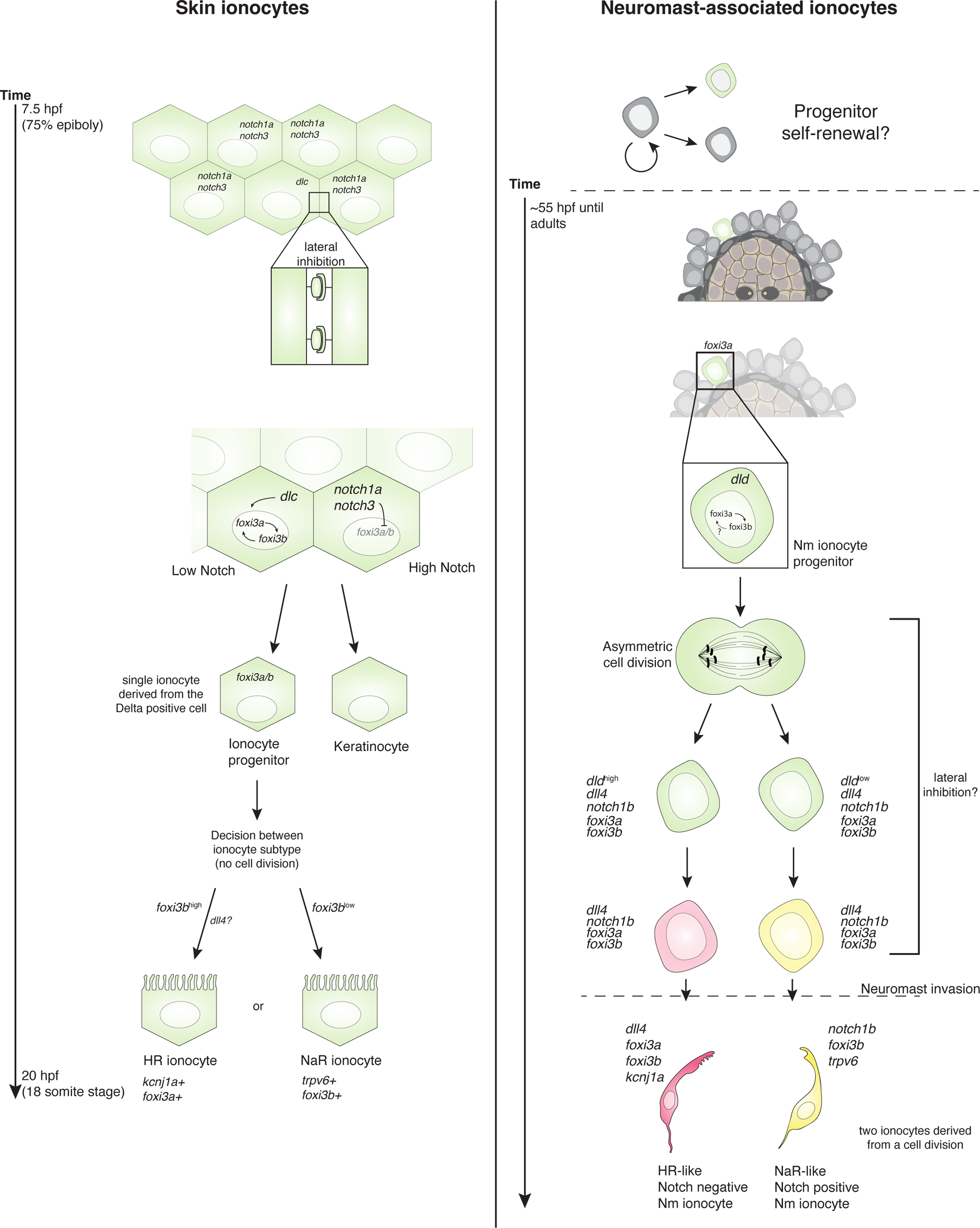
Comparison of ionocyte differentiation in the skin and the neuromasts. Model of tissue-specific ionocyte differentiation in zebrafish. Skin ionocytes are derived from *dlc* positive cells, and one ionocyte progenitor is formed without a cell division. Skin ionocyte progenitors then become either a NaR or HR ionocyte based on the expression of *foxi3b* and/or *dll4* (based on results from ^20,21,93^. Nm ionocytes, on the other hand, are derived from a cell division and give rise to a Nm ionocyte pair. The resulting two cells consist of one Notch-positive (*notch1b*) and one Notch-negative (*dll4*) cell, and sustained Notch signaling controls their survival. It is known that progenitors are derived from *krtt1c19e*-positive skin cells, but how the self-renewal of this single-cell progenitor takes place is not understood.

## Videos

**Supplementary Video 1: Nm ionocytes are derived from a cell division.** Neuromast in a 3 dpf *Tg(−8.0cldnB:lyn-EGFP)^zf106Tg^* zebrafish larva. A Nm ionocyte progenitor is seen dividing and the resulting pair invades the neuromast. Scale bar = 10 µm.

**Supplementary Video 2: *dld* is upregulated prior to progenitor cell division.** Time-lapse of a *dld:H2B-EGFP;ubi:H2A-mCherry* 3 dpf transgenic zebrafish shows *dld* is upregulated during cytokinesis. Scale bar = 10 µm. Gamma was changed in the *dld:H2B-EGFP* channel to allow for visualization of dim structures.

**Supplementary Video 3: *Nm ionocytes in a 4 dpf dll4 sibling*.** Time-lapse of a *dld:H2B-EGFP;cldnb:H2A-mCherry* transgenic larva shows a mature Nm ionocyte pair over the course of 32 hours. Scale bar = 10 µm. Gamma was changed in the *dld:H2B-EGFP* channel to allow for visualization of dim structures.

**Supplementary Video 4: *Cell death of Nm ionocytes is observed in 4 dpf dll4 mutants*.** Time-lapse of a *dll4* mutant larva showing a pair of Nm ionocytes, labeled by *dld:H2B-EGFP*, that migrate into a neuromast and die shortly after differentiation. Scale bar = 10 µm. Gamma was changed in the *dld:H2B-EGFP* channel to allow for visualization of dim structures.

## Notes

### Competing Interest Statement

The authors have declared no competing interest.

